# Peptide Information Compression Technology (PICT): A New Frontier in Therapeutic Peptide Discovery

**DOI:** 10.1101/2024.12.10.627447

**Authors:** Hong Wang, Zhu-Yin Wang, Wei-Ming Jiang, Qin-Qin Wang, Lin Miao, Deng Wang, Hai-Tao Wang, Shiu-Feng Tung, Jin-Rui Yang, Bin Yang, Xu-Bo Liang, Zhen-Wan Li, Xue-Ping Wang, Sheng Zou, Feng He, Xiao-Tao Guo, Yuan Yan, Kai-Min Xu, Xiao-Qi Wang, Xin-Bo Wang, Hui-Qing Liu, Xiang-Qun Li

**Affiliations:** Zonsen PepLib Biotech Inc., Zhuzhou, Hunan, China, 412000; Shenzhen PeptiBoom Investment Co., Ltd,Shenzhen, Guangdong, China, 518122

**Keywords:** Therapeutic peptides, Peptide Information Compression Technology (PICT), pentapeptide library, high-throughput screening (HTS), TSLP/TSLPR/IL7Rα inhibitors, MC3R antagonists

## Abstract

Therapeutic peptides hold tremendous market potential, yet the discovery methods for active peptides lag considerably behind those for small molecules and biological macromolecule drugs. Due to their respective limitations, traditional peptide compound synthesis and display libraries fail to meet the growing demands of the peptide drug market. Here, we introduce a novel Peptide Information Compression Technology (PICT) and, based on this technology, a large-scale physical peptide library and its accompanying high-throughput screening (HTS) platform. By screening for inhibitors of TSLP/TSLPR/IL7Rα and antagonists of MC3R at both the molecular and cellular levels, we have identified active molecules in the single-digital micromolar and nanomolar ranges, respectively. This demonstrates the immense application value of Peptide Information Compression Technology in the efficient discovery of peptide drugs.

## 1. Introduction

The field of peptide drug discovery has undergone a significant evolution, starting from the early 20th century with the therapeutic use of insulin, a naturally occurring peptide, marking a milestone in the history of peptide drugs^1,2^. This marked the beginning of understanding peptides’ therapeutic potential, leading to an expanded interest in synthetic peptides as drugs. As novel therapeutic molecules and excellent drug carriers, peptides have been widely used in immunotherapy, diabetes, obesity, cancer therapy, immune diseases, endocrine disorders, metabolic & cardiovascular disease, antiviral & antimicrobial therapy, acromegaly, etc^3,4^.

Peptide drugs offer unique advantages over small molecules and larger biologicals like monoclonal antibodies, including high specificity and potency, lower toxicity, biocompatibility, and synthetic accessibility. Despite their significant potential, the discovery and development of therapeutic peptide drugs encounter several bottlenecks, such as stability and degradation issues, poor oral bioavailability, low membrane permeability, and delivery challenges. Techniques have been developed to overcome these drawbacks, such as peptide cyclization (head-to-tail cyclization^5^, stapled peptides^6,7^, etc.), backbone modification^8,9^, termini protection^10,11^, lipidation^12,13^, PEGylation^14–16^, and conjugation to plasma proteins^17,18^ to solve the instability problem. The membrane penetration properties of peptide drugs can also be improved with the help of cell-permeable peptides (CPPs)^19,20^, receptor-mediated endocytosis^21,22^, and backbone modifications^23,24^.

Current methods in peptide drug discovery primarily involve automated solid-phase peptide synthesis (SPPS)^25–27^, liquid-phase peptide synthesis (LPPS)^28,29^, and HTS of synthetic libraries^30–32^. However, these methods often face limitations in diversity and complexity. Developing new peptide libraries is a response to these limitations, aiming to create more diverse, structurally complex, and modified peptides. These new libraries incorporate non-natural amino acids, cyclic peptides, and other modifications, enhancing stability and bioavailability and targeting a broader range of biological targets.

The drive to develop new peptide libraries stems from the need to overcome the intrinsic limitations of current peptide drugs and to explore novel therapeutic targets. These libraries are designed to enhance the pharmacokinetic properties, increase the stability and bioavailability of peptide drugs, and enable the targeting of previously ’undruggable’ biological pathways. Several methods are available for screening peptide drug candidates, such as phage display technology^33–35^, mRNA display^36–39^, one-bead one-compound (OBOC) technology^40–42^, Affinity Selection Mass Spectrometry (AS-MS)^43–46^, peptide microarray^47,48^, virtual screening^49–52^, etc.

Here, we introduce a new technique for discovering peptide drug candidates termed Peptide Information Compression Technology (PICT) and its applications using 2 case studies as examples. It ingeniously uses a small number of sizeable cyclic peptide molecules (>50aa) to display a vast array of peptide sequence combinations, thereby compressing peptide sequence information of tens of millions or even billions into tens of thousands of sizeable cyclic peptide molecules. Due to the unique structure of cyclic peptides, a cyclic peptide can construct a library very efficiently, containing a wide range of peptide sequences. For example, within one 80-mer cyclic peptide, there can be 80 different sequences of dipeptides, tripeptides, tetrapeptides, pentapeptides, and so on, up to 80-mer peptides (Fig. 1A). This structure demonstrates up to 6,320 unique peptide sequences within a single molecule, achieving an impressive compression ratio 1:6320 for embedded amino acid sequence information. This method significantly streamlines the process of building peptide libraries, makes obtaining highly active peptide molecules directly through HTS possible, reduces the workload, and increases efficiency in peptide drug discovery (Fig. 1B).

**Fig. 1.**
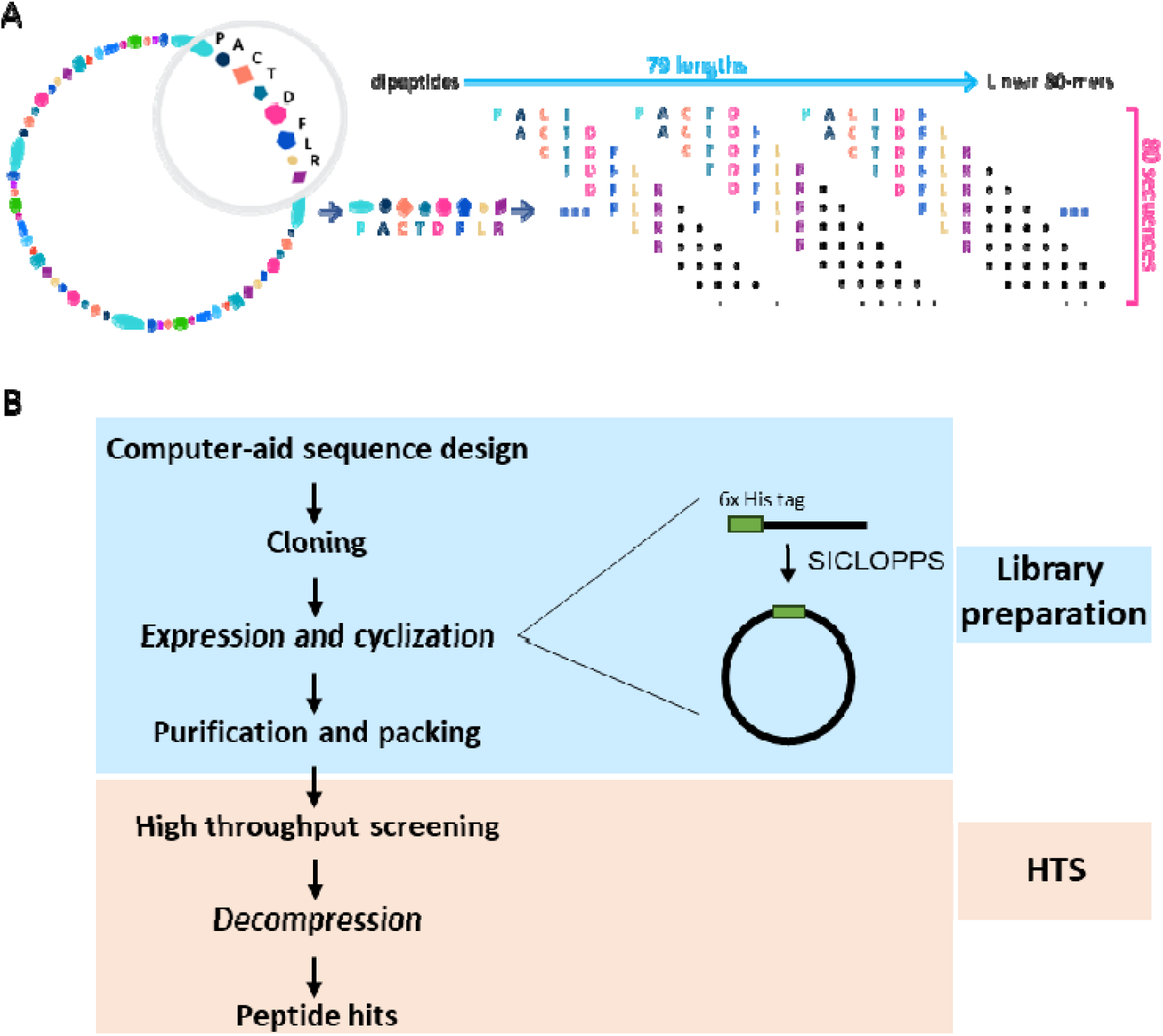
Design and preparation of the library. (A) Schematic diagram of PICT technology. Any region of the 80-mer cyclic peptide can be expanded into a linear peptide, then divided into 80 linear dipeptides, 80 linear tripeptides, 80 linear tetrapeptides, and so on, up to 80 linear 80-peptides. The compression ratio of peptide information is about 1:6000. B) Construction and screening process of the ZSenithFive® peptide library. 1. Design the sequence of 80-mer cyclic peptides through computer-aided methods; 2. Construct the genes encoding the 80-mer cyclic peptides into an expression vector used for peptide cyclization; 3. Express the constructed plasmids in *E. coli.* and perform intracellular cyclization; 4. Purify the 80-mer cyclic peptides on a large scale and store them in aliquots; 5. Use the ZSenithFive® peptide library to conduct HTS against designated target proteins at biochemical and cellular levels; 6. Decompress the active molecules obtained from HTS to yield shorter active peptides; 7. After multi-dimensional activity confirmation, the decompressed peptides can be used as candidates for subsequent preclinical development.

## 2. Results

### 2.1 Characteristics of the complete pentapeptide library

Based on PICT technology, we have successfully constructed an 80-mer cyclic peptide library containing all possible pentapeptides, termed “complete pentapeptide library” or “ZSenithFive® library.” We meticulously designed each sequence of the 80-mer cyclic peptides in the library, ensuring diversity and structural flexibility. Each 80-mer cyclic peptide contains a YCFHHHHHH sequence, essential for cyclization and purification. The remaining 71 positions are randomly distributed among the 20 amino acids, with no significant differences in the proportion of different amino acids (Fig. 2A). The average isoelectric point (pI) of the entire ZSenithFive® library is about 8.08, indicating slight alkalinity, with most (>75%) of the sequences having a pI in the range of 6-10. Moreover, under physiological conditions at pH 7.4, the average net charge is +1.38, slightly positive, enhancing the peptides’ usability and the universality of the screening methods. Additionally, we conducted secondary structure predictions for all sequences, calculating the proportion of helices, sheets, and disordered regions in each sequence. The results showed that the average percentages of helices, sheets, and disordered structural elements are 49.79%, 32.76%, and 18.46% (Fig. 2C), indicating that most of the peptide sequences are flexible yet capable of forming a small amount of secondary and even tertiary structures (Fig. 2D). This makes them suitable for target proteins with relatively complex ligand-binding pockets.

**Fig. 2.**
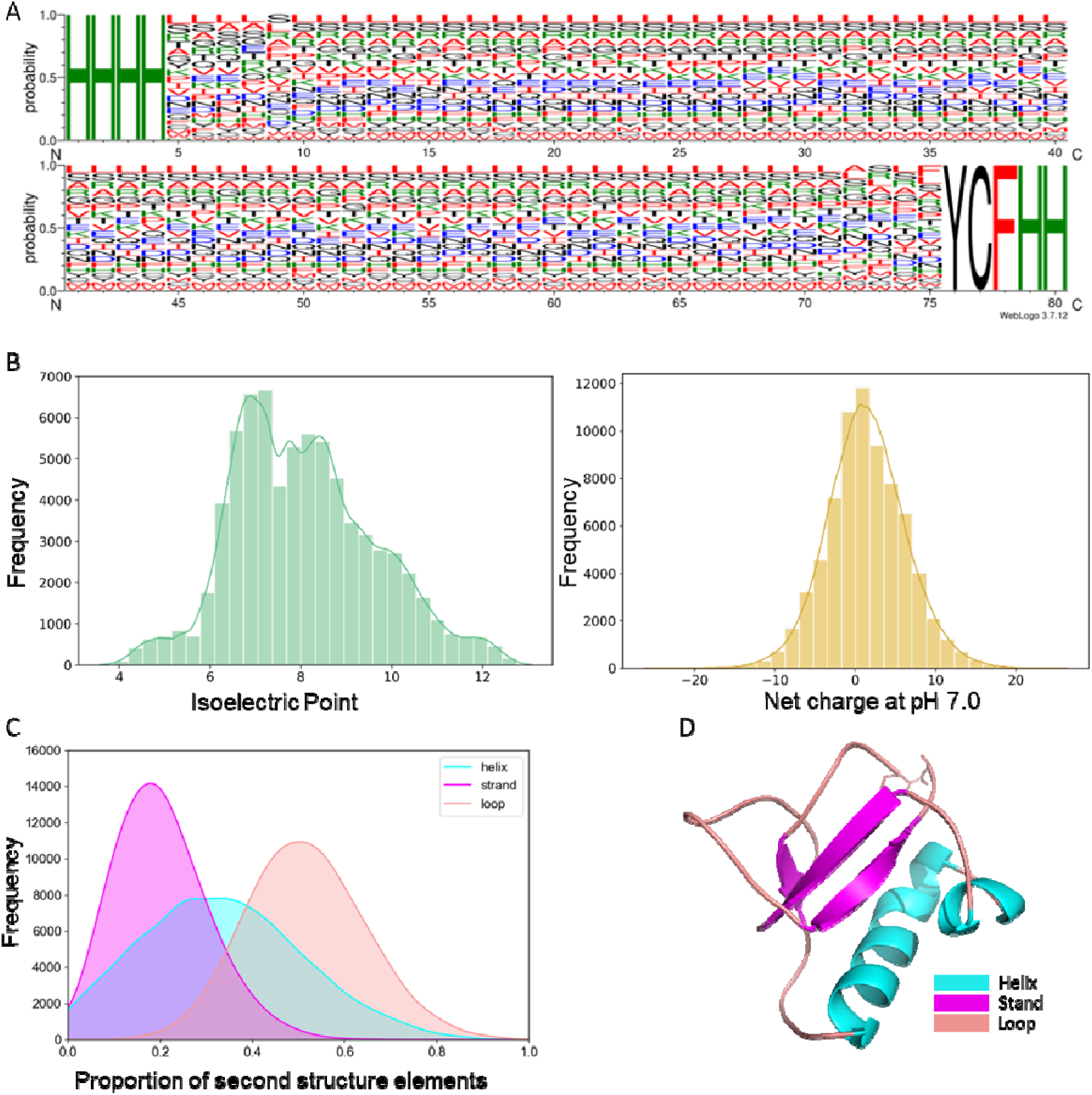
Characterization of the ZSenithFive® peptide library. (A) Amino acid composition chart for each position of the 72,960 80-mer cyclic peptide sequences. Each amino acid is represented by a single letter and differentiated by color: polar amino acids (black), basic amino acids (green), acidic amino acids (blue), and hydrophobic amino acids (red), with the letter size indicating the proportion of that amino acid at that position. (B) Predicted isoelectric points and electrostatic charge distribution at pH 7.0 for all 80-mer cyclic peptides. Both histograms specify the number of bins as 30, with curves representing density curves that describe the distribution shape of the variables. (C) Distribution of predicted secondary structural elements across all 80-mer cyclic peptide sequences. (D) Schematic illustration of potential tertiary structures within the 80-mer cyclic peptides. The color for secondary structural elements is consistent with Fig. C.

### 2.2 Application of the ZSenithFive® library for HTS

Using split-intein circular ligation of peptides and proteins (SICLOPPS) methods^53^, 72,960 80-mer peptides were expressed and cyclized in *E.coli* bacteria, isolated, and validated with HPLC-MS to guarantee their purity was higher than 90%. Then, they were sub-packaged into seven hundred and sixty 96-well plates in a "one-well-one-peptide" format and stored at -20 L. Like other synthetic libraries, our library can be directly used for high-throughput molecular and cellular screening.

#### 2.2.1 Inhibitor discovery of TSLP/TSLPR/IL7R**α** complex

TSLP is a cytokine that plays a crucial role in initiating and persisting allergic inflammation, making it a compelling target for therapeutic intervention. TSLP has been identified as a critical mediator in various allergic conditions, including asthma, atopic dermatitis, and allergic rhinitis. It acts upstream in the allergic cascade, influencing dendritic cells to induce a Th2-type immune response. Targeting TSLP offers the potential to modulate this response early, thereby attenuating the allergic reaction^54–56^. The focus on developing peptide-based drugs targeting TSLP revolves around designing peptides that can either inhibit the interaction of TSLP with its receptor or modulate the downstream signaling pathways^57–59^. The specificity and high affinity of peptides make them suitable candidates for disrupting these protein-protein interactions.

Time-resolved Fluorescence Energy Transfer (TR-FRET) is a practical combination of Time-Resolved Fluorescence (TRF) and Förster Resonance Energy Transfer (FRET). It combines the low background characteristic of TRF with the homogeneous detection form of FRET, offering higher throughput, fewer false positives/negatives, and improved flexibility, reliability, and sensitivity^60^. Consequently, we repackaged 72,960 sequences of 80-mer cyclic peptides into 456 384-well plates and conducted HTS using the TR-FRET technology (Fig S4A). We calculated and compared the Z and Z’ values for all plates (Fig. 3A), which are used to assess the suitability of HTS conditions and the quality of the test data and to track changes in experimental quality. The average Z and Z’ values of the 456 plates were 0.57 and 0.67, respectively, indicating good separation between the sample and reference signals in the detection system, with most plates meeting HTS standards. Plates with Z values or Z’ values less than 0 were excluded from analysis and selection.

**Fig. 3.**
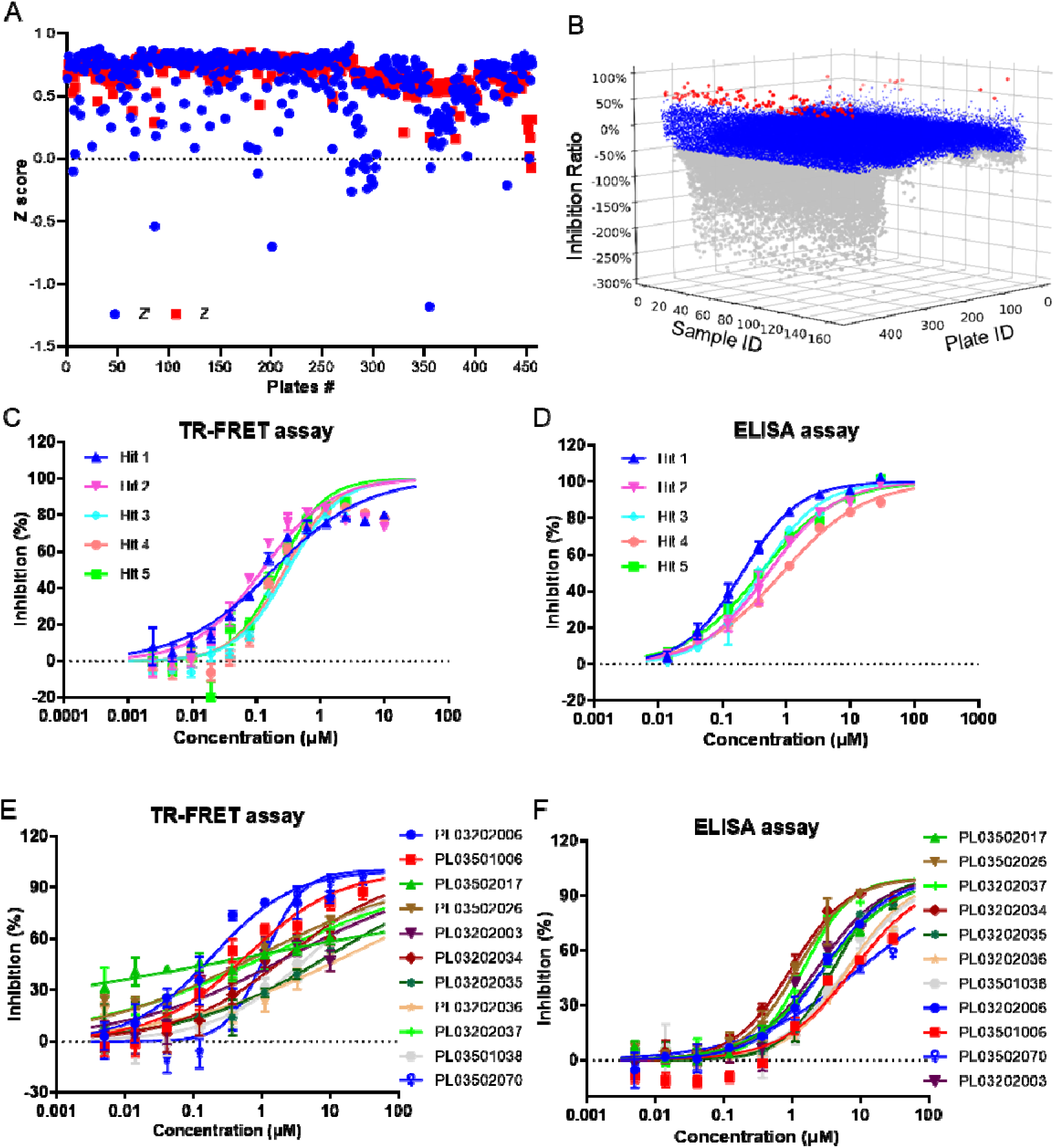
High-throughput screening and decompression for TSLP/TSLPR/IL7Ra inhibitors. (A) Z-value statistics for each 384-well plate after HTS. Red represents Z values, and blue represents Z’ values, with the vast majority of Z and Z’ values greater than 0.5, indicating the data quality meets HTS standards. (B) The inhibition percentage of each 80-mer cyclic peptide sample was converted and displayed in 3D form. 72,960 80-mer cyclic peptides were aliquoted into 456 384-well plates, with 160 samples per plate. Inhibition rates below -50% are shown in gray, greater than or equal to -50%, but less than 50% are in blue, and rates above 50% are in red. (C) Concentration-dependent inhibition curves for five representative 80-mer cyclic peptides measured using the TR-FRET method show significant inhibitory effects as the sample concentration increases for Hits 1 to 5. (D) Concentration-dependent inhibition curves for five representative 80-mer cyclic peptides measured using the ELISA method show significant inhibitory effects as the sample concentration increases for Hits 1 to 5. (E) Concentration-dependent inhibition curves for 11 decompressed short peptides were measured using the TR-FRET method, with some samples showing significant inhibitory effects as the sample concentration increased. (F) Concentration-dependent inhibition curves for 11 decompressed short peptides were measured using the ELISA method, with some samples showing significant inhibitory effects as the sample concentration increased.

Moreover, we calculated the inhibition rate of each sample on the binding of TSLP and IL-7Rα &TSLPR. Most samples had inhibition rates between -50 % and 50%, and 132 sequences showed inhibition rates greater than 50%, from which 86 hits were selected for the second round of multiple single-concentration re-screenings (Fig. 3B). Simultaneously, we developed an ELISA detection method^61^ for cross-validation of active molecules obtained from TR-FRET HTS. After re-screening, 20 hits consistently showed inhibitory effects on TSLP/TSLPR/IL7Rα binding at different concentrations. Then, these hits underwent IC_50_ testing in four duplicate wells in three independent repeat experiments. The IC_50_ values of these 20 samples, tested using the ELISA method, ranged from 0.61 to 8.69 μM, while the IC_50_ values tested using the TR-FRET method ranged from 0.07 to 6.42 μM (Table S1). Their inhibitory curves were shown in Fig. 3C and Fig. S6A for the TR-FRET assay and Fig. 3D and Fig. S6B for the ELISA assay, respectively. We analyzed the sequences of these 20 80-mer peptides and selected Hit2, Hit7, and Hit12 for decompression.

At the outset of designing the 80-mer peptides, we had already planned a decompression strategy for the PICT technology. We applied this to the decompression design and screening of Hit2, Hit7, and Hit12. During the decompression process, for each 80-mer hit, we designed 40 cyclic peptides of 30 amino acids and 40 corresponding linear peptides, 240 peptides in total, which were expressed and cyclized in *E. coli.* Then, the same TR-FRET method was used to screen these decompressed peptides for their ability to inhibit the interaction between TSLP and IL-7Rα &TSLPR. After the decompression screening, we obtained a series of 30-amino acid linear and cyclic peptides with inhibitory activity. By cross-validating with ELISA and TR-FRET, we identified peptides with IC_50_ values of 0.98-8.02 μM under ELISA conditions and 0.19-17.63 μM under TR-FRET conditions (Fig. 3E, 3F, and Table S2). PL03202037 and PL03202006 exhibited good inhibitory effects on TSLP/TSLPR/IL7Rα binding in both detection systems. Except for PL03502070, a cyclic peptide, all other peptides inhibiting the binding of TSLP and IL-7Rα &TSLPR were linear peptides. Finally, through HTS and decompression, we obtained several peptides from the library that can inhibit TSLP/IL7Rα/TSLPR at digital micromolar levels.

#### 2.2.2 Antagonist of Melanocortin Receptor 3

The research into peptide drug discovery targeting the Melanocortin 3 Receptor (MC3R) represents a significant area of interest in metabolic disorders and obesity management. MC3R, a G protein-coupled receptor (GPCR) in the melanocortin receptor family, regulates energy homeostasis, food intake, and metabolism. The melanocortin system, particularly MC3R and MC4R, is integral in controlling energy balance^62,63^. MC3R’s role in modulating energy homeostasis and metabolic processes makes it a target for treating obesity and related metabolic disorders^64,65^. Research in peptide drug discovery for MC3R primarily revolves around developing selective agonists or antagonists. Agonists aim to stimulate MC3R activity, potentially reducing food intake and increasing energy expenditure^66^. Antagonists, conversely, can help elucidate the physiological role of MC3R by blocking its function^67^. More interestingly, it has been reported recently that genetic deletion or pharmacological inhibition of MC3R may improve the dose responsiveness to Glucagon-like peptide 1 (GLP1) agonists^68^.

As we know, MC3R can signal through both the Gs/i G-protein/cAMP pathway and the Gq G-protein/calcium pathway to regulate various cellular functions. The binding of an agonist activates Gq downstream of MC3R, leading to an increase in intracellular calcium concentration. Intracellular calcium concentration can rapidly increase from 100-200nM to approximately 100μM, allowing for the measurement of transient changes in cytosolic calcium concentration using fluorescent calcium dyes. Before commencing HTS, as a preliminary step, the concentration-dependent response of ACTH and NDP-α-MSH-induced calcium flux in MC3R cells was assessed using the FLIPR Calcium 5 assay kit, as illustrated in Fig. S6A. This assay utilized a stable CHO-K1/Gα15/MC3R cell line that overexpressed Gα15 and MC3R. The EC_80_ values for ACTH and NDP-α-MSH-induced calcium flux in MC3R cells were 0.89 μM and 0.76 μM, respectively, as depicted in Fig. S6B. Subsequently, the IC_50_ of SHU9119 for inhibiting MC3R cells was determined at the selected ACTH and NDP-α-MSH concentrations (Fig. S6C). Finally, we chose ACTH as the agonist to stimulate CHO-K1/Gα15/MC3R cells for HTS of MC3R antagonists.

We used the validated HTS system described above to conduct HTS with 72,960 80-cyclic peptides from the ZSenithFive® peptide library. The peptide test concentration was set at 2 μM, and each peptide was tested in duplicates. In the ACTH control group, 384-well plates with a Z factor less than 0.4 were excluded from the data analysis phase (Fig. 4A). After calculating the agonistic and inhibitory activities for each sample, we selected 90 samples for a second round of single-concentration rescreening (Fig. 4B). After rescreening, five 80-cyclic peptides with inhibition rates greater than 50% were chosen for multi-concentration testing to determine their IC_50_ values. Each sample was tested in duplicate wells across two independent experiments. The results showed that the IC_50_ values for these five candidate molecules inhibiting the MC3R receptor were 0.28±0.07 μM, 0.60±0.18 μM, 0.54±0.06 μM, 0.41±0.04 μM, and 0.59±0.16 μM respectively (Fig. 4C). Finally, due to its highest inhibitory activity, Peptide hit 1 was selected for further decompression.

**Fig. 4.**
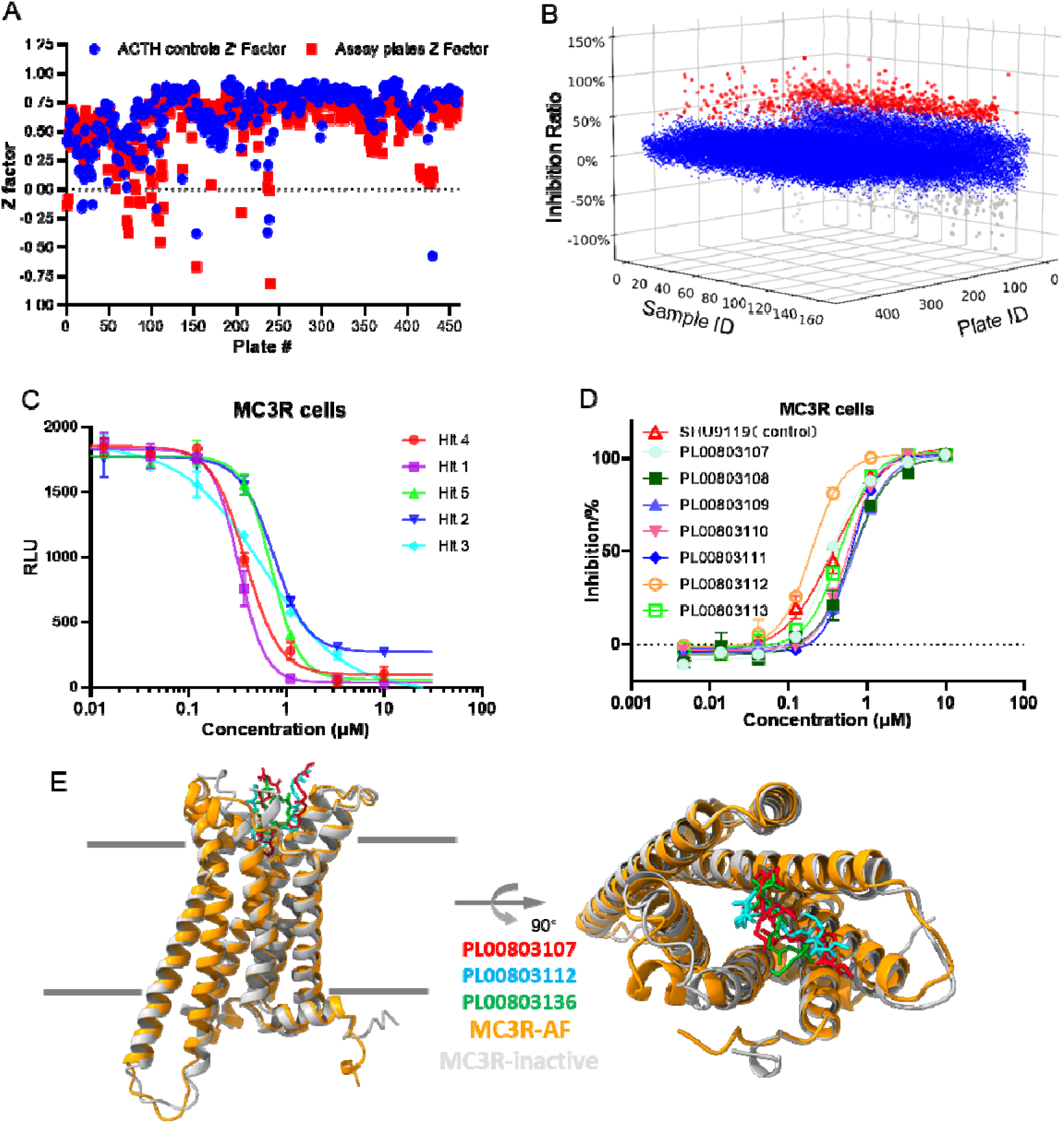
High-throughput screening and decompression for MC3R antagonists. (A) Z-value statistics for each 384-well plate after HTS. Red indicates Z values, and blue indicates Z’ values, with most Z and Z’ values exceeding 0.5, indicating that the data quality meets HTS standards. (B) Each 80-mer cyclic peptide sample’s inhibition percentage is converted and displayed in 3D form. 72,960 80-mer cyclic peptides were aliquoted into 456 384-well plates, with 160 samples per plate. Inhibition rates below -50% are shown in gray, greater than or equal to -50% but less than 50% in blue, and above 50% in red. (C) Concentration-dependent inhibition curves for five representative 80-mer cyclic peptides on MC3R cells, with Hits 1 to 5 showing significant inhibitory effects as the sample concentration increases. (D) Concentration-dependent inhibition curves for seven representative decompressed short peptides on MC3R cells show significant inhibitory effects as the sample concentration increases. Some samples exhibited activity superior to the positive control SHU9119. E) Schematic representation of the complex structures between three active peptides and the MC3R receptor predicted by AlphaFold. In the predicted complex structures, the conformation of the MC3R receptor (orange, MC3R-AF) is highly similar to the inactive conformation of MC3R in GPCRdb.

We adopted a decompression strategy similar to that used for the TSLP target and conducted multiple decompression rounds on Hit1. In the first round of decompression, we designed 40 cyclic peptides, each consisting of 40 amino acids and 40 corresponding linear peptides, making a total of 80 peptides. We used HTS methods to assess their ability to inhibit ACTH-induced activation of the MC3R receptor. We obtained a series of 40-amino acid linear peptides with inhibitory activity at a concentration of 2μM. We selected some peptides and tested them in a 2-fold serial dilution from 10μM across eight concentrations to verify if they exhibit concentration-dependent inhibitory activity. The results confirmed that these peptides exhibited significant inhibitory effects, with IC_50_ values ranging from 0.3 μM to 1.0 μM (Table S5).

Sequence analysis of these inhibitory peptides led to the second round of decompression, during which we designed several 20-amino acid cyclic and corresponding linear peptides. Among these, the cyclic peptide PL00803083 and the linear peptide PL00803082 showed the best activity. As shown in Fig. S9, PL00803083 had an IC_50_ of approximately 2 μM, while PL00803082 had an IC_50_ of roughly 13.5 μM. With the shortening of the peptide sequence, the cyclic peptides exhibited better inhibitory activity than their 40-amino acid linear counterparts.

Compared to SHU9119, the length of PL00803083 is still relatively longer. Thus, we conducted the third and fourth decompression rounds, designing a range of 9 to 19 amino acid cyclic peptides. These peptides were then tested using the above methods to determine their ability to inhibit ACTH-induced MC3R activation. Several peptides had inhibitory activity close to or even higher than SHU9119 (Fig.4D and S10). PL00803112 was the most potent, having an IC_50_ of 0.26±0.08 μM, twice as inhibitory as SHU9119 (IC_50=_ 0.5±0.16 μM). PL00803107, PL00803108, PL00803109, PL00803110, PL00803111, PL00803113, PL00803135, and PL00803136 had activities close to that of SHU9119. PL00803134, PL00803137, PL00803138, and PL00803139 showed weaker activity than SHU9119 (Table S6). The complex structures of active peptides (PL00803107, PL00803112, and PL00803136) with MC3R, as predicted by AlphaFold2, show that all three most active peptides bind to the extracellular ligand-binding pocket of MC3R. After binding with the peptides, the receptor’s conformation closely resembles the inactive conformation reported in GPCRdb (Fig. 4E).

#### 2.2.3 Selectivity of Hits for Melanocortin Receptors

To investigate whether the MC3R inhibitory molecules we obtained exhibit selectivity towards other melanocortin receptors, we selected three of the most potent MC3R antagonists, PL00803112, PL00803107, and PL00803136, for inhibition activity testing on CHO-K1 cells overexpressing other melanocortin receptors (MC2R, MC4R, and MC5R) and Gα15 protein. The testing method was consistent with that used for inhibiting the MC3R receptor, except that the agonist was changed to NDP-α-MSH. As evidenced by their IC_50_ values, the inhibitory activity of the three active molecules against NDP-α-MSH differed slightly from that against ACTH (Fig. 5A). The positive control SHU9119 had an IC_50_ value of 0.14 μM for inhibiting NDP-α-MSH-induced MC4R activation, while PL00803112, PL00803107, and PL00803136 had IC_50_ values of 79.55 μM, 54.29 μM, and 40.62 μM, respectively (Fig. 5B). SHU9119 exhibited no selectivity between MC3R and MC4R. The inhibitory activity of these three peptides against MC4R was weak, indicating strong selectivity. Notably, PL00803112 showed 1600 times greater inhibitory activity against MC3R than against MC4R, while PL00803107 and PL00803136 showed 10 to 50 times higher inhibitory activity against MC3R than MC4R. The peptides we screened displayed selectivity for MC3R. Additionally, for MC2R and MC5R, the inhibitory activity of the three peptides was significantly lower than that against MC3R, but the selectivity was not as pronounced as for MC4R (Fig. 5C and 5D). Through screening the ZSenithFive® peptide library and decompression, we identified a series of peptides with selective inhibitory activity against the MC3R receptor. Among them, PL00803112 showed the most vigorous activity, reaching 0.04 μM, surpassing the activity of SHU9119, and exhibiting 1600 times selectivity for MC4R.

**Fig. 5.**
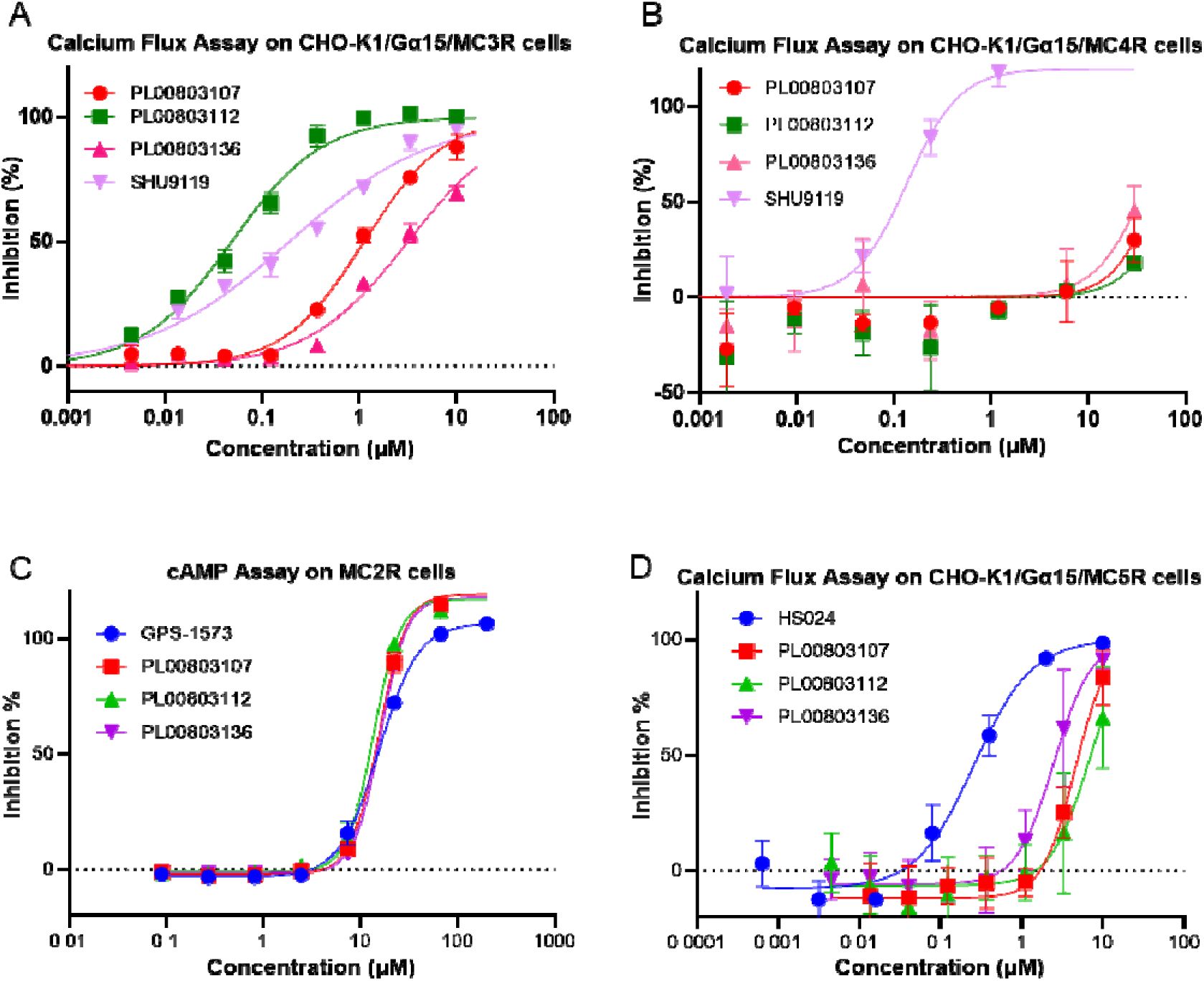
Comparison of the inhibitory activity of MC3R candidate antagonists on other Melanocortin receptors. A) Three active molecules showed significant inhibitory activity against MC3R in the Calcium Flux assay, with PL00803112 exhibiting superior activity to the positive control SHU9119. B) The three active molecules demonstrated poor inhibitory activity against MC4R in the Calcium Flux assay, significantly lower than the positive control SHU9119. C) The three active molecules showed poor inhibitory activity against MC5R in the Calcium Flux assay, significantly lower than the positive control HS024. D) The three active molecules exhibited significant inhibitory activity against MC2R in the cAMP assay, comparable to the activity of the positive control GPS-1573.

**Fig. 6.**
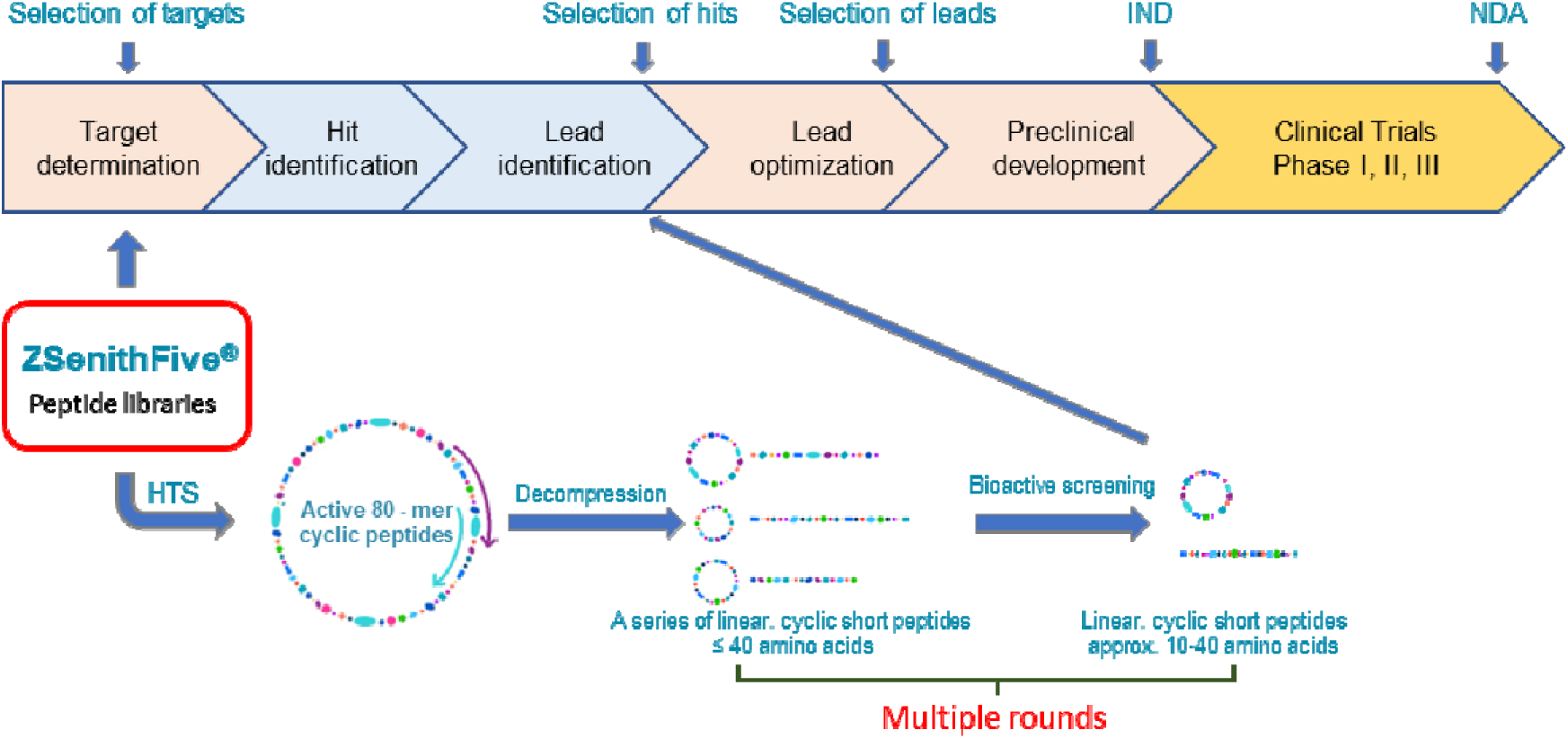
Schematic diagram of the high-throughput screening and decompression process for the ZSenithFive® peptide library. The ZSenithFive® peptide library can be used for HTS of various target types. After obtaining active 80-mer peptide molecules in the initial screening, 3-4 rounds of stepwise decompression are conducted to reduce the peptide length to 10-40 amino acids, thereby obtaining lead compounds. These then enter the lead optimization and preclinical development phase, ultimately advancing to Investigational New Drug (IND) applications for clinical trials.

## 3. Discussion

Through peptide information compression technology (PICT), we have built a peptide library containing 72,960 80-mer cyclic peptides, which not only covers all sequence information of di-, tri-, tetra-, and pentapeptide but also harbors other sequence information from 6-mer up to 80-mer peptides, totally nearly 500 million unique sequences.

By conducting HTS against various targets, 80-mer peptide hit compounds can be easily and effectively obtained. Subsequent multiple decompression rounds allowed us to discover lead peptides with suitable length and high potency for further optimization, as shown in our screening comparing against TSLP/TSLPR/IL7Rα and MC3R. These and other screening data (unpublished) fully demonstrate the value of the complete pentapeptide library.

Compared to peptide libraries built from other technologies, such as phage display and mRNA display, which are typically mixtures of different compounds, our ZSenithFive® library demonstrates many advantages, to list a few:

1. Each peptide is individually expressed, purified, quantified, and QC verified.
2. Compound stability is higher because of the peptide-bond cyclic structure.
3. The library is intrinsically complete for sequences of pentapeptides and below due to the design strategy.
4. Peptides are packaged in 96-well HTS format for the convenience of the subsequent screening.
5. Rebuilding the entire library is easy and cheap with the biosynthesis process.
6. Biological assay methods for hit screening are not limited to affinity/binding assays. Cell-based functional assays, even in vivo assays, can also be chosen.
7. There is no interference between compounds during the screening process. Hence, we believe the false positive or false negative rate during the hit screening is lower.

Despite these advantages, our library may also have some potential flaws: the flexibility of the cyclic peptide structure might lead to non-natural folding due to cyclization; some peptides might form higher-order structures that cannot interact with target proteins in screening assays; issues of non-specific binding in peptide-protein interactions; and costs and time associated with decompression, etc. However, solutions can be found to address these issues, such as peptide microarray technology, which can significantly accelerate screening times and reduce the amount of target protein required, complementary peptide libraries that can supplement the structural and sequence diversity of the 80-mer cyclic peptides, and the assistance of computers and artificial intelligence which can shorten the time for decompression, reduce the number of short peptides that need to be synthesized during decompression, and increase the success rate of decompression, etc.

## 4. Methods

### 4.1 Preparation of the pentapeptide library

The pentapeptide library sequences were designed and compressed into approximately 80,000 80-mer peptides, with about 60% derived from natural genomic sequences of organisms like *E. coli, Saccharomyces cerevisiae, Arabidopsis thaliana, Homo sapiens,* and *Mus musculus*. The remaining sequences were manually designed. Genomic DNA for the natural sequences was extracted using a homemade kit, and primers were designed following standard Gibson assembly protocols. PCR amplification used Phusion High-Fidelity DNA Polymerases, and PCR fragments were purified. Codon-optimized sequences for E. coli were chemically synthesized and assembled into double-stranded fragments via polymerase cycling assembly (PCA). These sequences were cloned into a pET15b vector with Npu DnaE intein, linearized at a NheI restriction site, and inserted using Gibson Assembly^®^. Plasmid constructs were verified by DNA sequencing and expressed in ArcticExpress (DE3) cells, then analyzed by Tricine-SDS-PAGE.

Cells were grown in an auto-induction medium for large-scale expression, lysed, and peptides purified using Nickel Magbeads. The molecular weight and purity of the peptides were assessed by MALDI-TOF MS and HPLC. Peptides meeting purity criteria (>90%) were quantified, aliquoted into 96-well plates, and lyophilized. The resulting “ZSenithFive peptide library” contains 72,960 unique 80-mer cyclic peptides stored in 760 vacuum-packed 96-well plates at -20°C for high-throughput screening.

### 4.2 HTS of “ZSenithFive peptide library” for TSLP and IL-7R**α**/TSLPR antagonist

On screening day, peptides were dissolved in sterile water, transferred to 384-well plates, and diluted using Automated Liquid Handlers (Nimbus96, Hamilton). TR-FRET was applied to screen peptides inhibiting the protein-protein interaction (PPI) between TSLP and IL-7Rα/TSLPR. The assay measured the interaction by the fluorescence ratio emitted by Allophycocyanin (665 nm) to Europium (620 nm). Initial tests with varying concentrations of Biotin-TSLP (16 nM, 8 nM, 4 nM, 2 nM, 1 nM) and IL-7Rα/TSLPR (16 nM, 8 nM, 4 nM, 2 nM, 1 nM) determined optimal concentrations for the HTS system, with 1 nM each showing the best signal-to-background ratio. The anti-TSLP antibody, used as a positive inhibitor, showed a concentration-dependent inhibition with an IC50 of approximately 0.6 μg/ml (∼ 4 nM). For validation, two peptides from the library were tested, yielding a Z factor greater than 0.5, suitable for HTS. In a 384-well plate, peptide, Biotin-TSLP, IL-7Rα/TSLPR, fluorescent donor (SA-Eu, Europium-Streptavidin), and acceptor (Anti-hFc-APC, Allophycocyanin-conjugated AffiniPure Goat Anti-Human IgG, Fc Fragment Specific) were mixed, incubated at room temperature for 2 hours, and read for TR-FRET signals using a fluorescent plate reader (Cytation5, Biotek).

For ELISA assay, IL-7Rα/TSLPR were diluted with coating buffer (0.05 mol/L carbonate buffer pH 9.6) to different concentrations (0.125 μg/ml, 0.25 μg/ml, 0.5 μg/ml, 1 μg/ml, 2 μg/ml, 4 μg/ml) and coated with 25μl/well overnight at 4°C. The next day, wash 3∼5 times with washing buffer (0.05% Tween-20 in TBS, pH 7.4). Add blocking solution (20mg/ml BSA in TBS, pH 7.4) and incubate at 37°C for blocking. After 3∼5 times washing, add an appropriate amount of Biotin-TSLP protein and different concentrations of Anti-TSLP antibody (30 μg/ml, then 2-fold serial dilution of 8∼10 concentrations), and incubate at 37°C for 1 hour. Wash off the unbound fraction, add SA-HRP, and incubate at 37°C for 1 hour. Wash subsequently, add TMB for color development, terminate the reaction with 1M HCL, and read OD_450_ in a plate reader (Cytation5, Biotek).

The primary screening of the ZSenithFive peptide library at 2 μM was followed by hit confirmation via TR-FRET and ELISA at multiple doses to calculate IC50 values. In the TR-FRET assay, the emission ratio (ER) was calculated by dividing the acceptor emission value (665 nm) by the donor emission value (620 nm).

The peptide inhibition rate was calculated using the equation:

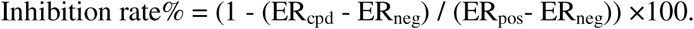

The positive signal control Z′ factor was calculated using the equation:

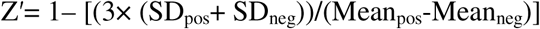

The antagonist screening plate Z factor was calculated using the equation:

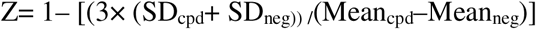

In which ‘cpd’ means compound, ‘pos’ means positive control, and ‘neg’ means negative control.

Selected 80-mer peptide hits were further decompressed and screened with FRET and ELSA as described above. Data was fitted to a four-parameter logistic model to determine IC50 values using GraphPad Prism version 10.0.0 for Windows.

### 4.3 HTS of “ZSenithFive peptide library” for Melanocortin receptor binders

For the FLIPR assay, the ZSenithFive peptide library was stored and dissolved as previously described above. Chinese hamster ovary cells stably expressing human MC3R (CHO-K1/MC3R/Gα15) were cultured following the manufacturer’s instructions in Nutrient Mixture F-12 Ham, supplemented with 10% fetal bovine serum, 1× penicillin/streptomycin, 100 μg/ml hygromycin, and 200 μg/ml Zeocin in a humidified atmosphere (90%-95%) with 5% CO2 at 37°C, and passaged every 2-3 days or when 80%-90% confluent. 8000-10000 cells were seeded into a 384-well black clear-bottom plate and incubated overnight. Calcium 5 dye was diluted 20-fold with a specific buffer (20 mM HEPES, pH 7.4, 1.26 mM CaCl_2_, 0.49 mM MgCl_2_, 0.41 mM MgSO_4_, 5.37 mM KCl, 0.44 mM KH_2_PO_4_, 0.34 mM Na_2_HPO_4_, 136.9 mM NaCl, 4.2 mM NaHCO_3_, and 5.6 mM D-Glucose). After discarding the cell media, 50 μl of Calcium 5 dye solution was added per well and incubated in the dark for 2 hours. Agonists and antagonists were diluted and added to a 384-well compound plate. The cell and compound plates were transferred to a Fluorescent Imaging Plate Reader (FLIPR Tetra) to measure calcium ion changes induced by ACTH or NDP-α-MSH in MC3R cells, with readings taken every second for 160 seconds. SHU9119’s inhibition of calcium ion changes was also detected over 320 seconds, with inhibition calculated from signals recorded between 170-320 seconds. For subsequent screenings, 0.9 μM ACTH and 1.5 μM SHU9119 were used as the agonist and antagonist, respectively, representing EC80 and IC90 values. Peptides were dissolved to 200 μM, then diluted to 10 μM using Automated Liquid Handlers. Cells were seeded into 384-well plates, incubated overnight, treated with Calcium 5 dye, and assayed for peptide activity using the FLIPR Tetra reader at a final peptide concentration of 2 μM.

All the FLIPR data was collected in ScreenWorks version 4.2. The mean 1 relative fluorescence units (RFU) of 1s to 10s, the maximum 1 RFU of 11 to 160s, the mean 2 RFU of 161 to170s point time, and the maximum 2 RFU of 171 to 320s statistic were exported to Excel.

For agonist analysis, △RFU1= RFU_maximum1_-RFU_mean1_.

For antagonist analysis, △RFU2= RFU_maximum2_-RFU_mean2_.

The percentage of activation and inhibition was calculated as the following formulation:

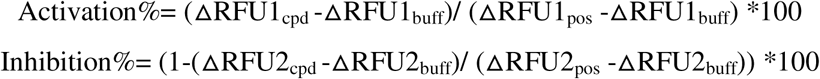

The positive control Z′ factor was calculated using the equation:

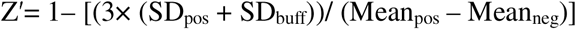

The agonist screening plate Z factor was calculated using the equation:

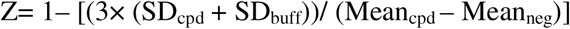

The antagonist screening plate Z factor was calculated using the equation:

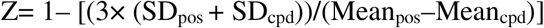

In this case, ‘cpd’ meant compound, ‘pos’ meant positive control, and ‘buff’ meant buffer control. The data was fitted to a four-parameter logistic model in GraphPad Prism version 10.0.0 for Windows to calculate EC50 or IC50 values.80-mer peptide hits were selected according to the IC_50_ and decompressed up to 4 rounds. Decompression peptides were screened and confirmed by FLPR assay as described above.

### 4.4 Structure prediction and analysis

The sequence log of 72,960 80-mers was generated and visualized using WebLogo 3.7.12^69^. The secondary structure of all 72,960 80-mers was predicted using PSSpred (Protein Secondary Structure prediction), a simple neural network training algorithm for accurate protein secondary structure prediction^70^. The percentage of different secondary structures H, E, C (H: alpha-helix, E: extended strand, C: the rest) in each sequence was calculated using Python pandas 2.1.4 and visualized in Microsoft Office Excel. The isoelectric points and net charges at pH 7.4 of 80-mer cyclic peptides were calculated with Python Peptides 0.3.2 and visualized in Microsoft Office Excel. The structure of PL00803107, PL00803112, and PL00803136 with MC3R were predicted with AlphaFold 2^71^ and visualized in Open-Source PyMOL.

## Supporting information

Supplemental Figures and Tabals

## Acknowledgments

Thanks to Ning Zhu, Fen Xie, and Hua Dai, who contributed significantly to this work but are not included in the author list.

## **5.** Author contributions

Project initiation and supervision: XQL, XBW, and HQL. Library design: ZYW, WMJ and QQW. Library construction: QQW, LM, XBL, SZ, FH, BY, XPW, XTG, KMX, HTW, and

YY. Decompression design: QQW. High-throughput screening: JRY, ZWL, and DW. Data interpretation: DW, HW, and SFT. Figure preparation: HW, DW, HTW, and XQW. Manuscript writing: HW, WJ, and DW. All authors read and contributed to the manuscript.

## **6.** Data and materials availability

All data are available in the main text or supplementary materials.

## Reference

1 Muttenthaler, M., King, G. F., Adams, D. J. & Alewood, P. F. Trends in peptide drug discovery. Nat Rev Drug Discov 20, 309–325, doi:10.1038/s41573-020-00135-8 (2021).

2 Scott, D. A. & Best, C. H. The Preparation of Insulin. Industrial & Engineering Chemistry 17, 238–240, doi:10.1021/ie50183a004 (1925).

3 Sharma, K., Sharma, K. K., Sharma, A. & Jain, R. Peptide-based drug discovery: Current status and recent advances. Drug Discov Today 28, 103464, doi:10.1016/j.drudis.2022.103464 (2023).

4 Wang, L. et al. Therapeutic peptides: current applications and future directions. Signal Transduct Target Ther 7, 48, doi:10.1038/s41392-022-00904-4 (2022).

5 Slough, D. P., McHugh, S. M. & Lin, Y. S. Understanding and designing head-to-tail cyclic peptides. Biopolymers 109, e23113, doi:10.1002/bip.23113 (2018).

6 Moiola, M., Memeo, M. G. & Quadrelli, P. Stapled Peptides-A Useful Improvement for Peptide-Based Drugs. Molecules 24, doi:10.3390/molecules24203654 (2019).

7 Ali, A. M., Atmaj, J., Van Oosterwijk, N., Groves, M. R. & Domling, A. Stapled Peptides Inhibitors: A New Window for Target Drug Discovery. Comput Struct Biotechnol J 17, 263–281, doi:10.1016/j.csbj.2019.01.012 (2019).

8 Ayikpoe, R. S. & van der Donk, W. A. Peptide backbone modifications in lanthipeptides. Methods Enzymol 656, 573–621, doi:10.1016/bs.mie.2021.04.012

9. (2021).

9 Ahn, J. M., Boyle, N. A., MacDonald, M. T. & Janda, K. D. Peptidomimetics and peptide backbone modifications. Mini Rev Med Chem 2, 463–473, doi:10.2174/1389557023405828 (2002).

10 Lucana, M. C. et al. Protease-Resistant Peptides for Targeting and Intracellular Delivery of Therapeutics. Pharmaceutics 13, doi:10.3390/pharmaceutics13122065 (2021).

11 Chandrashekar, C., Okamoto, R., Izumi, M. & Kajihara, Y. Chemical Modification of the N Termini of Unprotected Peptides for Semisynthesis of Modified Proteins by Utilizing a Hydrophilic Protecting Group. Chemistry 25, 10197–10203, doi:10.1002/chem.201901778 (2019).

12 Kowalczyk, R., Harris, P. W. R., Williams, G. M., Yang, S. H. & Brimble, M. A. Peptide Lipidation - A Synthetic Strategy to Afford Peptide Based Therapeutics. Adv Exp Med Biol 1030, 185–227, doi:10.1007/978-3-319-66095-0_9 (2017).

13 Zhang, L. & Bulaj, G. Converting peptides into drug leads by lipidation. Curr Med Chem 19, 1602–1618, doi:10.2174/092986712799945003 (2012).

14 Andrianov, A. K. Noncovalent PEGylation of protein and peptide therapeutics. Wiley Interdiscip Rev Nanomed Nanobiotechnol 15, e1897, doi:10.1002/wnan.1897 (2023).

15 Ambrosio, E. et al. Control of Peptide Aggregation and Fibrillation by Physical PEGylation. Biomacromolecules 19, 3958–3969, doi:10.1021/acs.biomac.8b00887 (2018).

16 Roberts, M. J., Bentley, M. D. & Harris, J. M. Chemistry for peptide and protein PEGylation. Adv Drug Deliv Rev 54, 459–476, doi:10.1016/s0169-409x(02)00022-4 (2002).

17 Wijesinghe, A., Kumari, S. & Booth, V. Conjugates for use in peptide therapeutics: A systematic review and meta-analysis. PLoS One 17, e0255753, doi:10.1371/journal.pone.0255753 (2022).

18 Bumbaca, B., Li, Z. & Shah, D. K. Pharmacokinetics of protein and peptide conjugates. Drug Metab Pharmacokinet 34, 42–54, doi:10.1016/j.dmpk.2018.11.001 (2019).

19 Xie, J. et al. Cell-Penetrating Peptides in Diagnosis and Treatment of Human Diseases: From Preclinical Research to Clinical Application. Front Pharmacol 11, 697, doi:10.3389/fphar.2020.00697 (2020).

20 Copolovici, D. M., Langel, K., Eriste, E. & Langel, U. Cell-penetrating peptides: design, synthesis, and applications. ACS Nano 8, 1972–1994, doi:10.1021/nn4057269 (2014).

21 Kato, Y. & Sugiyama, Y. Targeted delivery of peptides, proteins, and genes by receptor-mediated endocytosis. Crit Rev Ther Drug Carrier Syst 14, 287–331 (1997).

22 Habib, S. & Singh, M. Angiopep-2-Modified Nanoparticles for Brain-Directed Delivery of Therapeutics: A Review. Polymers (Basel) 14, doi:10.3390/polym14040712 (2022).

23 Barman, P. et al. Strategic Approaches to Improvise Peptide Drugs as Next Generation Therapeutics. Int J Pept Res Ther 29, 61, doi:10.1007/s10989-023-10524-3 (2023).

24 Li, X. et al. Backbone N-methylation of peptides: Advances in synthesis and applications in pharmaceutical drug development. Bioorg Chem 141, 106892, doi:10.1016/j.bioorg.2023.106892 (2023).

25 Coin, I., Beyermann, M. & Bienert, M. Solid-phase peptide synthesis: from standard procedures to the synthesis of difficult sequences. Nat Protoc 2, 3247–3256, doi:10.1038/nprot.2007.454 (2007).

26 Collins, J. M., Porter, K. A., Singh, S. K. & Vanier, G. S. High-efficiency solid phase peptide synthesis (HE-SPPS). Org Lett 16, 940–943, doi:10.1021/ol4036825 (2014).

27 Fields, G. B. Introduction to peptide synthesis. Curr Protoc Protein Sci Chapter 18, 18 11 11–18 11 19, doi:10.1002/0471140864.ps1801s26 (2002).

28 Sharma, A., Kumar, A., de la Torre, B. G. & Albericio, F. Liquid-Phase Peptide Synthesis (LPPS): A Third Wave for the Preparation of Peptides. Chem Rev 122, 13516–13546, doi:10.1021/acs.chemrev.2c00132 (2022).

29 Bayer, E. & Mutter, M. Liquid phase synthesis of peptides. Nature 237, 512–513, doi:10.1038/237512a0 (1972).

30 Quartararo, A. J. et al. Ultra-large chemical libraries for the discovery of high-affinity peptide binders. Nat Commun 11, 3183, doi:10.1038/s41467-020-16920-3 (2020).

31 Liu, R., Li, X., Xiao, W. & Lam, K. S. Tumor-targeting peptides from combinatorial libraries. Adv Drug Deliv Rev 110–111, 13-37, doi:10.1016/j.addr.2016.05.009 (2017).

32 Falciani, C., Lozzi, L., Pini, A. & Bracci, L. Bioactive peptides from libraries. Chem Biol 12, 417–426, doi:10.1016/j.chembiol.2005.02.009 (2005).

33 Bazan, J., Calkosinski, I. & Gamian, A. Phage display--a powerful technique for immunotherapy: 1. Introduction and potential of therapeutic applications. Hum Vaccin Immunother 8, 1817–1828, doi:10.4161/hv.21703 (2012).

34 Molek, P., Strukelj, B. & Bratkovic, T. Peptide phage display as a tool for drug discovery: targeting membrane receptors. Molecules 16, 857–887, doi:10.3390/molecules16010857 (2011).

35 Smith, G. P. Filamentous fusion phage: novel expression vectors that display cloned antigens on the virion surface. Science 228, 1315–1317, doi:10.1126/science.4001944 (1985).

36 Wilson, D. S., Keefe, A. D. & Szostak, J. W. The use of mRNA display to select high-affinity protein-binding peptides. Proc Natl Acad Sci U S A 98, 3750–3755, doi:10.1073/pnas.061028198 (2001).

37 Roberts, R. W. & Szostak, J. W. RNA-peptide fusions for the in vitro selection of peptides and proteins. Proc Natl Acad Sci U S A 94, 12297–12302, doi:10.1073/pnas.94.23.12297 (1997).

38 Newton, M. S., Cabezas-Perusse, Y., Tong, C. L. & Seelig, B. In Vitro Selection of Peptides and Proteins-Advantages of mRNA Display. ACS Synth Biol 9, 181–190, doi:10.1021/acssynbio.9b00419 (2020).

39 Lipovsek, D. & Pluckthun, A. In-vitro protein evolution by ribosome display and mRNA display. J Immunol Methods 290, 51–67, doi:10.1016/j.jim.2004.04.008 (2004).

40 Martinez-Ceron, M. C. et al. Latest Advances in OBOC Peptide Libraries. Improvements in Screening Strategies and Enlarging the Family From Linear to Cyclic Libraries. Curr Pharm Biotechnol 17, 449–457, doi:10.2174/1389201017666160114095553 (2016).

41 Lam, K. S. et al. Synthesis and screening of "one-bead one-compound" combinatorial peptide libraries. Methods Enzymol 369, 298–322, doi:10.1016/S0076-6879(03)69017-8 (2003).

42 Lam, K. S., Lebl, M. & Krchnak, V. The "One-Bead-One-Compound" Combinatorial Library Method. Chem Rev 97, 411–448, doi:10.1021/cr9600114 (1997).

43 van Breemen, R. B. & Muchiri, R. N. Affinity selection-mass spectrometry in the discovery of anti-SARS-CoV-2 compounds. Mass Spectrom Rev 43, 39–46, doi:10.1002/mas.21800 (2024).

44 Prudent, R., Lemoine, H., Walsh, J. & Roche, D. Affinity selection mass spectrometry speeding drug discovery. Drug Discov Today 28, 103760, doi:10.1016/j.drudis.2023.103760 (2023).

45 Muchiri, R. N. & van Breemen, R. B. Affinity selection-mass spectrometry for the discovery of pharmacologically active compounds from combinatorial libraries and natural products. J Mass Spectrom 56, e4647, doi:10.1002/jms.4647 (2021).

46 Jonker, N., Lingeman, H. & Irth, H. Direct Dynamic Protein-Affinity Selection Mass-Spectrometry. Chromatographia 72, 7–13, doi:10.1365/s10337-010-1586-x. (2010).

47 Szymczak, L. C., Kuo, H. Y. & Mrksich, M. Peptide Arrays: Development and Application. Anal Chem 90, 266–282, doi:10.1021/acs.analchem.7b04380 (2018).

48 Hundsberger, H. et al. Assembly and use of high-density recombinant peptide chips for large-scale ligand screening is a practical alternative to synthetic peptide libraries. BMC Genomics 18, 450, doi:10.1186/s12864-017-3814-3 (2017).

49 Tripathi, N. M. & Bandyopadhyay, A. High throughput virtual screening (HTVS) of peptide library: Technological advancement in ligand discovery. Eur J Med Chem 243, 114766, doi:10.1016/j.ejmech.2022.114766 (2022).

50 Amarasinghe, K. N. et al. Virtual Screening Expands the Non-Natural Amino Acid Palette for Peptide Optimization. J Chem Inf Model 62, 2999–3007, doi:10.1021/acs.jcim.2c00193 (2022).

51 Gahlawat, A. et al. Structure-Based Virtual Screening to Discover Potential Lead Molecules for the SARS-CoV-2 Main Protease. J Chem Inf Model 60, 5781–5793, doi:10.1021/acs.jcim.0c00546 (2020).

52 Kleandrova, V. V., Ruso, J. M., Speck-Planche, A. & Dias Soeiro Cordeiro, M.N. Enabling the Discovery and Virtual Screening of Potent and Safe Antimicrobial Peptides. Simultaneous Prediction of Antibacterial Activity and Cytotoxicity. ACS Comb Sci 18, 490–498, doi:10.1021/acscombsci.6b00063 (2016).

53 Osher, E. L. & Tavassoli, A. Intracellular Production of Cyclic Peptide Libraries with SICLOPPS. Methods Mol Biol 1495, 27–39, doi:10.1007/978-1-4939-6451-2_3 (2017).

54 Ebina-Shibuya, R. & Leonard, W. J. Role of thymic stromal lymphopoietin in allergy and beyond. Nat Rev Immunol 23, 24–37, doi:10.1038/s41577-022-00735-y (2023).

55 Adhikary, P. P., Tan, Z., Page, B. D. G. & Hedtrich, S. TSLP as druggable target - a silver-lining for atopic diseases? Pharmacol Ther 217, 107648, doi:10.1016/j.pharmthera.2020.107648 (2021).

56 Huston, D. P. & Liu, Y. J. Thymic stromal lymphopoietin:a potential therapeutic target for allergy and asthma. Curr Allergy Asthma Rep 6, 372–376, doi:10.1007/s11882-996-0006-7 (2006).

57 Mullard, A. FDA approves first-in-class TSLP-targeted antibody for severe asthma. Nat Rev Drug Discov 21, 89, doi:10.1038/d41573-022-00013-5 (2022).

58 Matera, M. G., Rogliani, P., Calzetta, L. & Cazzola, M. TSLP Inhibitors for Asthma: Current Status and Future Prospects. Drugs 80, 449–458, doi:10.1007/s40265-020-01273-4 (2020).

59 Park, S. et al. Synthesis and biological evaluation of peptide-derived TSLP inhibitors. Bioorg Med Chem Lett 27, 4710–4713, doi:10.1016/j.bmcl.2017.09.010 (2017).

60 Padros, J., Chatel, G. & Caron, M. Time-resolved Forster Resonance Energy Transfer Assays for Measurement of Endogenous Phosphorylated STAT Proteins in Human Cells. J Vis Exp, doi:10.3791/62915 (2021).

61 Tabatabaei, M. S. & Ahmed, M. Enzyme-Linked Immunosorbent Assay (ELISA). Methods Mol Biol 2508, 115–134, doi:10.1007/978-1-0716-2376-3_10 (2022).

62 Yanik, T. & Durhan, S. T. Specific Functions of Melanocortin 3 Receptor (MC3R). J Clin Res Pediatr Endocrinol 15, 1–6, doi:10.4274/jcrpe.galenos.2022.2022-5-21 (2023).

63 Cai, M. & Hruby, V. J. The Melanocortin Receptor System: A Target for Multiple Degenerative Diseases. Curr Protein Pept Sci 17, 488–496, doi:10.2174/1389203717666160226145330 (2016).

64 Sweeney, P. et al. The melanocortin-3 receptor is a pharmacological target for the regulation of anorexia. Sci Transl Med 13, doi:10.1126/scitranslmed.abd6434 (2021).

65 Olney, J. J., Navarro, M. & Thiele, T. E. Targeting central melanocortin receptors: a promising novel approach for treating alcohol abuse disorders. Front Neurosci 8, 128, doi:10.3389/fnins.2014.00128 (2014).

66 Moscowitz, A. E. et al. The Importance of Melanocortin Receptors and Their Agonists in Pulmonary Disease. Front Med (Lausanne) 6, 145, doi:10.3389/fmed.2019.00145 (2019).

67 Ericson, M. D. et al. Bench-top to clinical therapies: A review of melanocortin ligands from 1954 to 2016. Biochim Biophys Acta Mol Basis Dis 1863, 2414–2435, doi:10.1016/j.bbadis.2017.03.020 (2017).

68 Dahir, N. S. et al. Inhibition of the melanocortin-3 receptor (MC3R) causes generalized sensitization to anorectic agents. bioRxiv, doi:10.1101/2023.12.05.570114 (2023).

69 Crooks, G. E., Hon, G., Chandonia, J. M. & Brenner, S. E. WebLogo: a sequence logo generator. Genome Res 14, 1188–1190, doi:10.1101/gr.849004 (2004).

70 Yan, R., Xu, D., Yang, J., Walker, S. & Zhang, Y. A comparative assessment and analysis of 20 representative sequence alignment methods for protein structure prediction. Sci Rep 3, 2619, doi:10.1038/srep02619 (2013).

71 Jumper, J. et al. Highly accurate protein structure prediction with AlphaFold. Nature 596, 583–589, doi:10.1038/s41586-021-03819-2 (2021).

