## Supplemental Figures and Tabals for "Peptide Information Compression Technology (PICT): A New Frontier in Therapeutic Peptide Discovery"

**This PDF file includes:**

Materials

Supplementary Figs. S1 to S10

Table S1 to S7

**Materials and Methods**

**1.1 Cell line, proteins, plasmids, and reagents**

The melanocortin family overexpression cell line (CHO-K1/MC3R/Gα15, M00242; CHO-K1/MC4/Gα15, M00272; CHO-K1/MC2/Gα15, M00336; CHO-K1/MC5/Gα15, M00243) was purchased from Genscript (Nanjing, China); ArcticExpress (DE3) from Agilent(Santa Clara, CA, USA); Biotin-TSLP (His, Avi tag, TSP-H82Eb) and IL-7RΑ & TSLPR (Human IL-7R alpha & TSLPR Heterodimer Protein, Fc Tag & Fc Tag, MALS verified, ILR-H5255) were from ACROBiosystems (Newark, DE, USA); Anti-TSLP from Biolegend (San Diego, CA, USA); SA-Eu (Europium-Streptavidin) was from AAT Bioquest (Pleasanton, CA, USA); Anti-hFc-APC (Allophycocyanin-conjugated AffiniPure Goat Anti-Human IgG, Fc Fragment Specific) was from Jackson ImmunoResearch (West Grove, Pennsylvania, USA); SA-HRP (Streptavidin, horseradish peroxidase-conjugated), fetal bovine serum (FBS), Hygromycin B, Zeocin, Trypsin, 384-well FRET plate and 384-well black clear-bottom plate were all from Thermofisher (Waltham, MA, USA); 384-well ELISA plate from Greiner (Kremsmünster, Austria); Calcium 5 Assay Kit from Molecular Devices (San Jose, CA, USA); Nutrient Mixture F-12 Ham from Sigma (St. Louis, MO, USA); Peptides were custom synthesized by Genscript (Nanjing, China); Primers and Oligos from General Biosystems (Durham, NC, USA); Phusion High-Fidelity DNA Polymerase and Gibson Assembly® Master Mix were from NEB (Ipswich, MA, USA); Restriction enzymes were from Takara (Beijing, China); pET15b was from Novagen (Hong Kong, China); Tryptone and Yeast Extract from OXOID (Hampshire, United Kingdom); other chemical reagents mainly were from Sigma (St. Louis, MO, USA).

**Supplementary Figures S1 to S9**

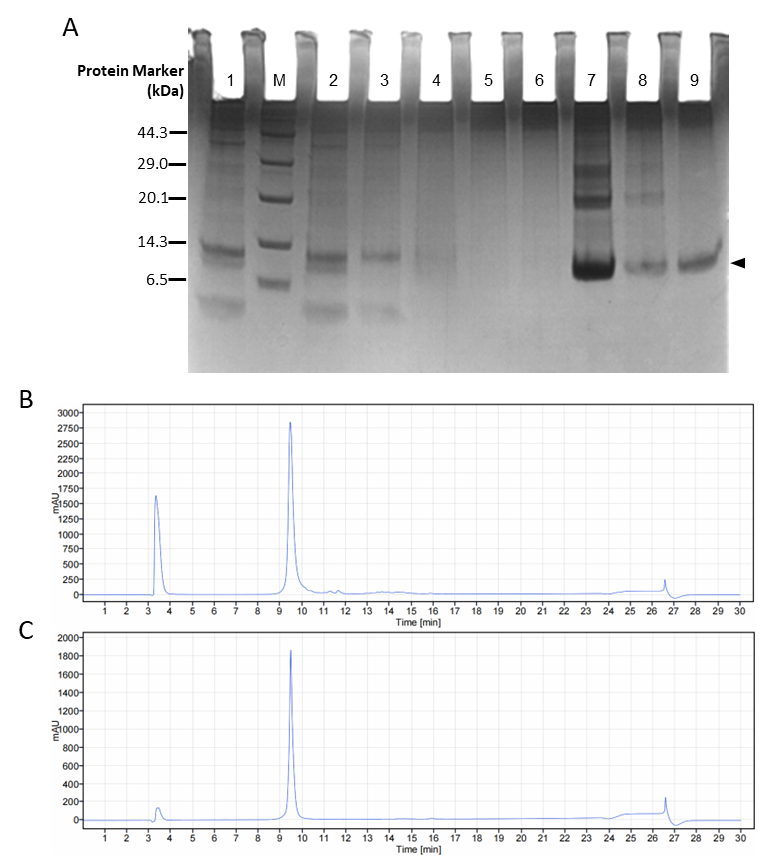

**Fig.S1.** Large-scale expression and purification of one 80-mer example. A) SDS-PAGE image of #527-59 peptide. The black triangle pointed to the target peptide. M: protein marker; 1: Overnight induced cells 2: Cell lysate; 3: Flow through of IMAC; 4: First washing of IMAC; 5: Second washing of IMAC; 6: First elution of IMAC 7: Second elution of IMAC; 8: Flow through of reversed-phase chromatography; 9: Elution of reversed-phase chromatography. B) HPLC curve of IMAC purified #527-59 peptide. C) HPLC curve of #527-59 peptide purified by IMAC plus reversed-phase chromatography. (IMAC: immobilized metal affinity of chromatography)

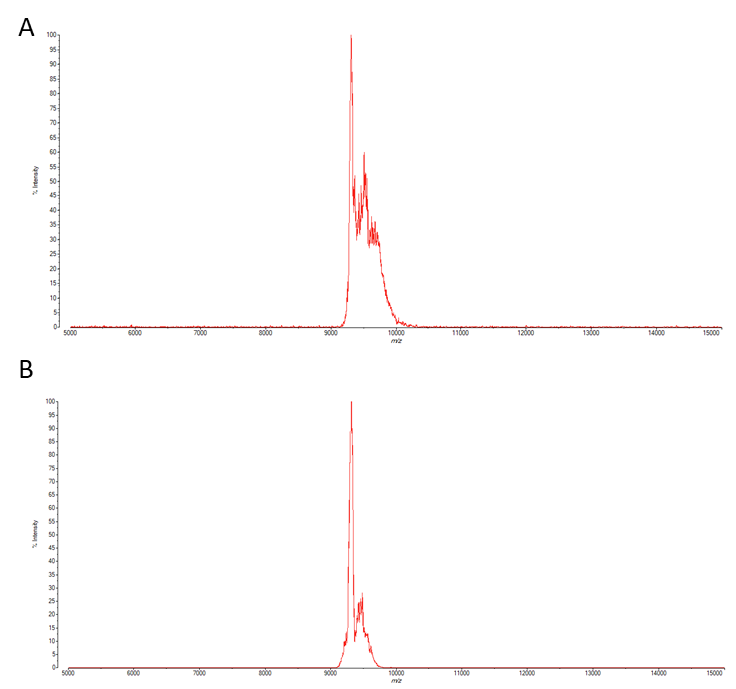

**Fig.S2.** MS report of purified #527-59 peptide. A) #527-59 peptide purified by IMAC. B) #527-59 peptide purified by IMAC plus reversed-phase chromatography.

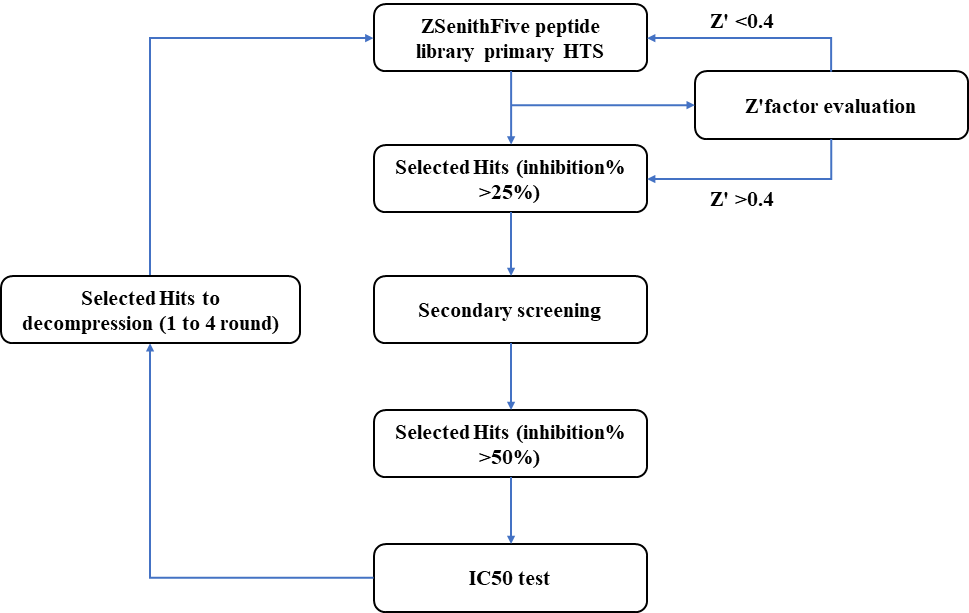

**Fig.S3.** High-throughput screening flowchart of the ZSenithFive peptide library. Firstly, establish a high-throughput screening method for different target proteins and screening goals and perform preliminary high-throughput screening using the ZSenithFive peptide library. Then, evaluate the initial screening using the Z’-factor. Samples with a Z’-factor less than 0.4 are retested, and those with a Z’-factor greater than 0.4 and an inhibition rate over 25% are selected as candidates for the second round of screening. Next, multi-concentration tests were performed on samples from the second round with inhibition rates greater than 50% to obtain their IC_50_ values. Finally, select the 3-5 sequences with the best IC_50_ values for 1-4 rounds of decompression, where the synthesized short peptides (linear or cyclic) are assessed for activity using the same method as the initial high-throughput screening (HTS).

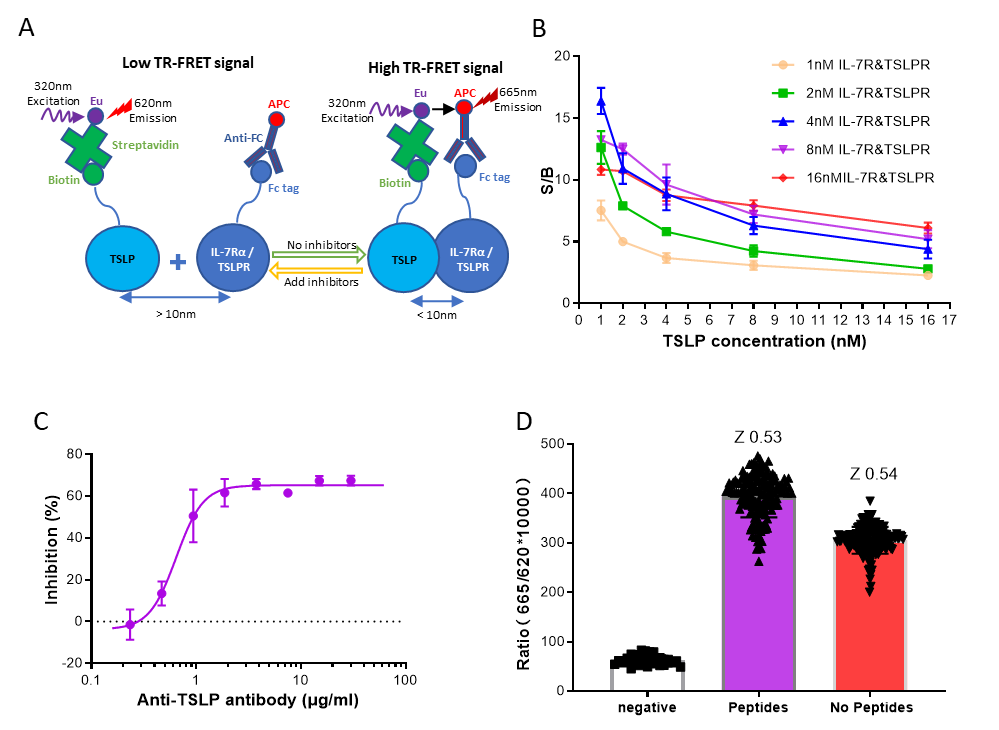

**Fig.S4.** Development and validation of TR-FRET assay for TSLP/TSLPR/ IL-7Rα. A) Schematic illustration of the TR-FRET assay. Streptavidin-Eu (SA-EU) coupled with biotinylated TSLP serves as the TR-FRET donor, and APC conjugated anti-Human Fc antibody coupled with Fc-IL7Rα/TSLPR serves as the acceptor. At the basal level, TSLP interacts with IL7Rα/TSLPR, yielding a high TR-FRET signal. The induced complex dissociation brings two fluorophores into further proximity upon treatment with inhibitors, generating a low TR-FRET signal. B) TR-FRET signal window value for the interaction between TSLP and IL-7Rα/TSLPR. TSLP and IL-7Rα/TSLPR were diluted to 16nM, 8nM, 4nM, 2nM, and 1nM for cross-combination. The S/B (signal-to-background) ratio, representing the TR-FRET signal of the two proteins at different concentrations relative to the blank group, is used to evaluate the system's window value. The maximum signal window can reach up to 15 times, indicating good screening applicability. Data are presented as mean ± SD (n=4). C) The dose-dependent inhibitory effect of TSLP antibodies on the binding of TSLP to IL-7Rα/TSLPR was tested with the TR-FRET assay. The anti-TSLP antibody was diluted to 30μg/ml and then serially diluted in 2-fold increments across eight concentrations. The maximum inhibitory effect of this anti-TSLP antibody did not reach 100%, but it showed a good concentration-dependent inhibitory effect with an IC_50_ of approximately 0.6μg/ml (4nM). Data are presented as mean ± SD (n=4). D) Validation of the TR-FRET method in a 384-well high-throughput screening system. In a 384-well plate, either with or without adding 5μM peptide, TSLP, IL-7Rα/TSLPR, and a pre-mixed SA-Eu and Anti-Human Fc-APC solution were sequentially added. After incubation, the TR-FRET signal was detected. We randomly selected 160 samples for testing. The Z value for the peptide test group was greater than 0.5, and the Z value for the group without peptide was greater than 0.5, with an S/B value greater than 5.

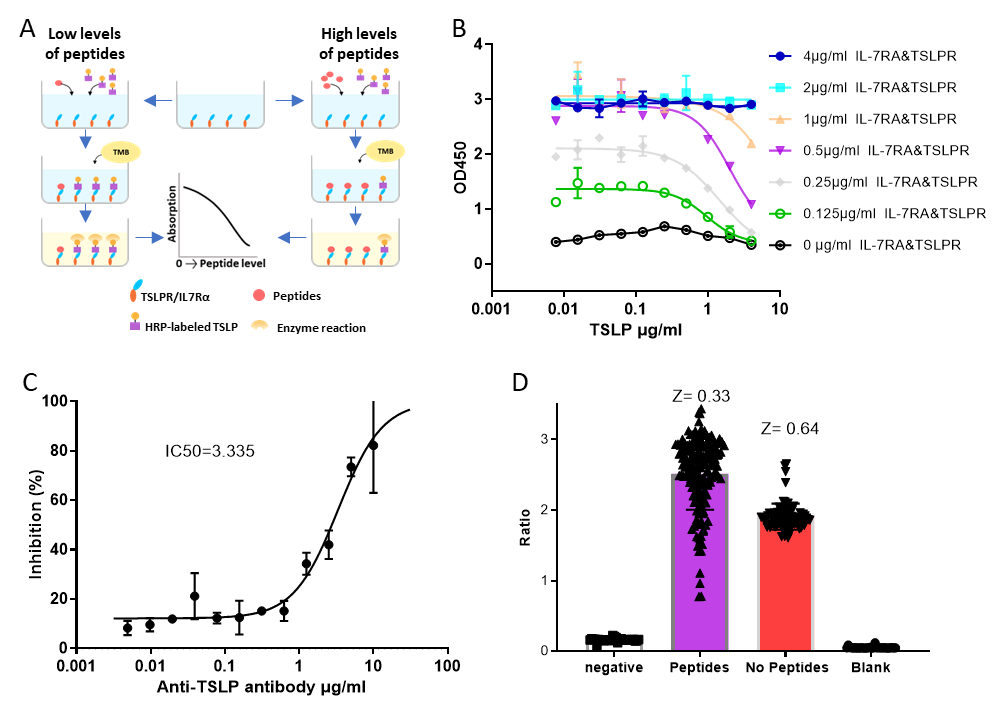

**Fig.S5.** Developing and validating ELISA assay for TSLP/TSLPR/ IL-7Rα. A. Schematic illustration of the competitive ELISA assay. The TSLPR/ IL-7Rα complex is immobilized on the surface of microplate wells and incubated with samples containing the TSLP and different levels of peptides. After the reaction, the activity of the microplate-bound enzyme is measured. B) The correlation between absorption and protein levels in samples was tested on a 384-well plate. Different concentrations of IL-7RΑ and TSLPR were coated. Then, pre-mixed samples of TSLP with SA-HRP were added, followed by incubation, washing, and color development. Coating with 0.5μg/ml of IL-7RΑ and TSLPR reached a plateau signal of around 3. Each concentration was set up in quadruplicate. Data are presented as mean ± SD (n=4). C) The dose-dependent inhibitory effect of TSLP antibodies on the binding of TSLP to IL-7Rα/TSLPR was tested with the ELISA assay. The anti-TSLP antibody was diluted to 10μg/ml and then serially diluted in 2-fold increments across 12 concentrations. This anti-TSLP antibody did not achieve 100% maximum inhibition, but it showed an excellent concentration-dependent inhibitory effect, with an IC_50_ of approximately 3.3 μg/ml (22nM). Each concentration was set up in quadruplicate. Data are presented as mean ± SD (n=4). D) Validation of the ELISA method in a 384-well high-throughput screening system was conducted. IL-7Rα/TSLPR was coated in a 384-well plate, followed by washing and blocking. Then, 5μM or no peptide was added, followed by the sequential addition of TSLP and SA-HRP pre-mix. After incubation, the OD_450_ signal was detected. We randomly selected 160 samples for testing. The Z value for the group without peptides was also greater than 0.5. The Z value for the peptide test group was slightly lower, at 0.33.

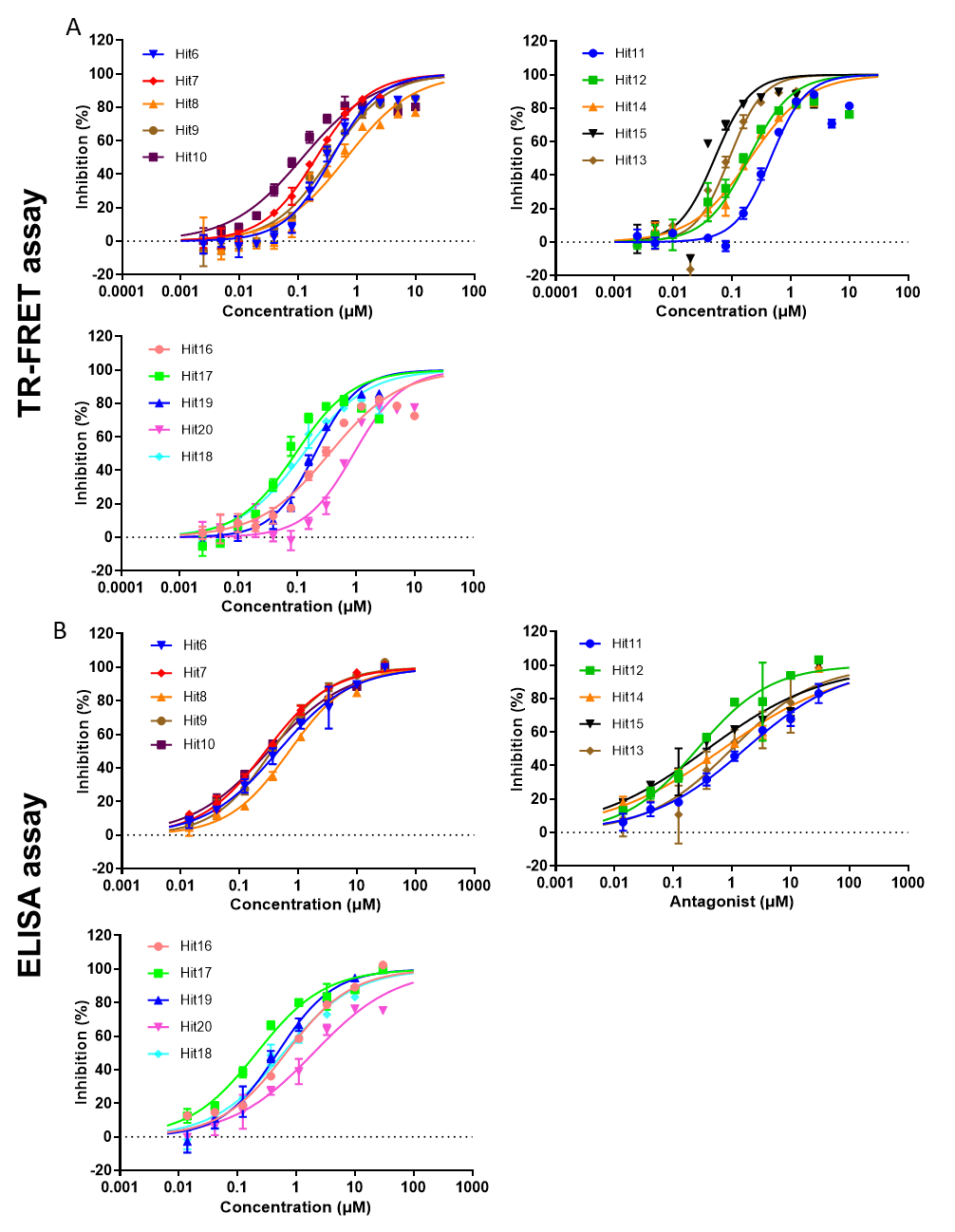

**Fig. S6.** The dose-dependent inhibitory effect of selected 80-mer hits on the binding of TSLP to IL-7Rα/TSLPR. A) Inhibitory curves of fifteen 80-mer hits with FR-FRET assay. B) Inhibitory curves of fifteen 80-mer hits with ELISA assay. The results of ELISA and TR-FRET were highly consistent.

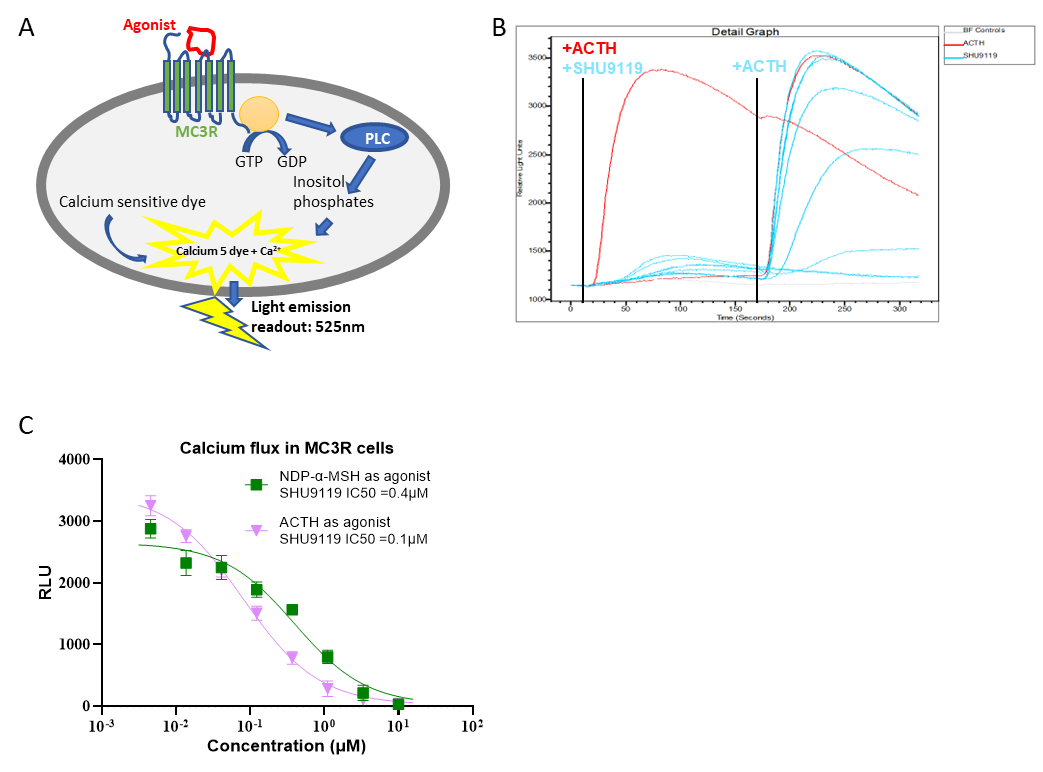

**Fig. S7** Development and validation of FLIPR assay for MC3R cells. A) Schematic illustration of the FLIPR assay. B) High-throughput screening curves simulated with ScreenWorks version 4.2 software. The red line represents the known agonist ACTH signal curve at 0.9μM, while the cyan lines represent the known inhibitor SHU9119 at different doses. Data from 0 to 160 seconds are for agonist screening; peptides that exhibit signals similar to the ACTH control group are selected as agonists. Data from 161 to 320 seconds are for inhibitor screening; peptides are added first and read for 160 seconds before ACTH is added to activate. If peptide samples in the wells show signal reduction similar to the SHU9119 control, they are selected as inhibitors. C) The IC_50_ of SHU9119 was 0.1μM and 0.4μM to 0.9μM ACTH and 0.76 μM NDP-α-MSH-induced calcium signals, respectively.

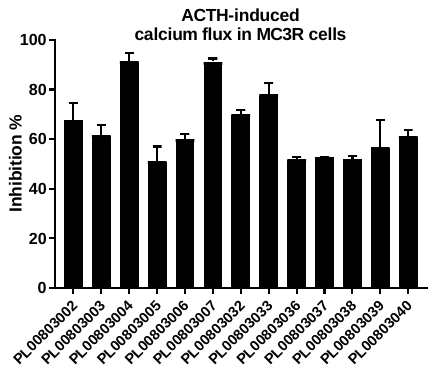

**Fig. S8.** Peptides yielded from the first round of decompression showed inhibitory activity against 0.9 μM ACTH-induced MC3R. The inhibition rate was obtained at a single concentration of 2μM.

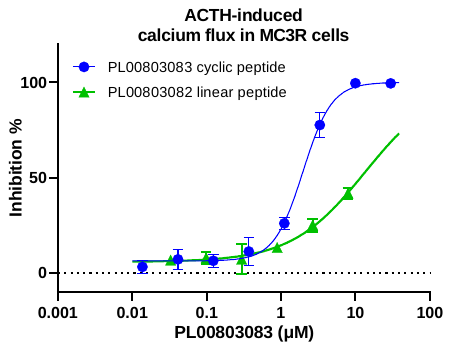

**Fig. S9.** Peptides yielded from the second round of decompression showed inhibitory activity against 0.9 μM ACTH-induced MC3R. PL00803083, a 20-amino acid cyclic peptide, had an IC_50_ of approximately 2μM for inhibiting ACTH-induced MC3R activation, while PL00803082, a 20-amino acid linear peptide, had an IC_50_ of roughly 13.5μM for the same inhibition.

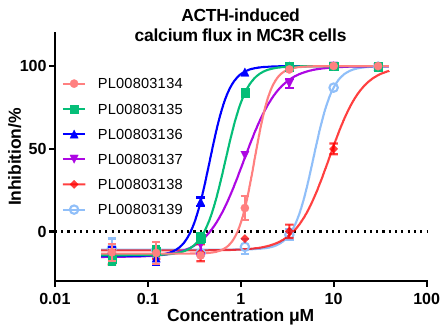

**Fig. S10.** Peptides from the fourth decompression round showed inhibitory activity against 0.9 μM ACTH-induced MC3R. Peptides designed in the fourth round had 7 to 12 amino acids. Among these, PL00803136 showed the most potent inhibition of 0.9 μM ACTH-induced MC3R activation, with an IC_50_ of 0.51±0.06 μM. All data were tested in two independent repeat experiments.

**Supplementary Table S1 to S7**

**Table S1**

IC_50_ values of 80-mer hits inhibiting the interaction between TSLP and IL-7Rα/TSLPR were obtained by ELISA and FRET assay. Average IC_50_ and standard deviation were calculated. 'NA' indicates that the IC_50_ was not tested.

| Hits ID | Library ID | IC_50_（μM）ELISA assay | | | | | IC_50_（μM）TR-FRET assay | | | | | |
| --- | --- | --- | --- | --- | --- | --- | --- | --- | --- | --- | --- | --- |
|  |  | test1 | test2 | test3 | Mean | SD | test1 | test2 | test3 | test4 | Mean | SD |
| Hit1 | 013-57 | 2.54 | 0.2 | 0.52 | 1.09 | 1.27 | 0.18 | 0.13 | 0.19 | 0.19 | 0.17 | 0.03 |
| Hit2 | 044-16 | 2.49 | 0.52 | 1.11 | 1.37 | 1.01 | 0.74 | 0.54 | 0.67 | 0.12 | 0.52 | 0.28 |
| Hit3 | 072-32 | 1.73 | 0.41 | 0.85 | 1.00 | 0.67 | 0.96 | 0.81 | 1.54 | 0.32 | 0.91 | 0.50 |
| Hit4 | 243-14 | 4.03 | 0.86 | 2.45 | 2.45 | 1.59 | 1.64 | 1.19 | 2.1 | 0.28 | 1.30 | 0.78 |
| Hit5 | 255-23 | 2.32 | 0.39 | 1.16 | 1.29 | 0.97 | 0.32 | 0.24 | 0.37 | 0.23 | 0.29 | 0.07 |
| Hit6 | 262-41 | 3.76 | 0.46 | 1.64 | 1.95 | 1.67 | 1.95 | 1.67 | 3.57 | 0.38 | 1.89 | 1.31 |
| Hit7 | 263-39 | 1.34 | 0.28 | 1.09 | 0.90 | 0.55 | 2.49 | 1.08 | 3.53 | 0.20 | 1.83 | 1.48 |
| Hit8 | 692-37 | NA | 0.71 | 1.66 | 1.19 | 0.83 | 6.83 | 5.35 | 12.81 | 0.67 | 6.42 | 5.01 |
| Hit9 | 695-25 | 1.58 | 0.36 | 0.7 | 0.88 | 0.63 | 5.53 | 2.68 | 7.71 | 0.36 | 4.07 | 3.22 |
| Hit10 | 704-18 | 2.06 | 0.32 | 0.86 | 1.08 | 0.89 | 3.34 | 6.41 | 6.34 | 0.12 | 4.05 | 2.99 |
| Hit11 | 714-76 | 16.78 | 1.69 | 5.13 | 7.87 | 7.91 | 6.14 | 0.76 | 0.58 | 0.46 | 1.98 | 2.77 |
| Hit12 | 715-93 | 1.19 | 0.26 | 0.68 | 0.71 | 0.47 | 1.23 | 1.06 | 1.6 | 0.19 | 1.02 | 0.60 |
| Hit13 | 720-26 | 21.57 | 1 | 3.49 | 8.69 | 11.23 | 0.59 | 0.46 | 0.23 | 0.09 | 0.34 | 0.22 |
| Hit14 | 730-46 | 11.48 | 0.7 | 5.28 | 5.82 | 5.41 | 0.23 | 0.29 | 0.21 | 0.20 | 0.23 | 0.04 |
| Hit15 | 743-62 | 2.86 | 0.39 | 2.75 | 2.00 | 1.40 | 0.09 | 0.08 | 0.06 | 0.05 | 0.07 | 0.02 |
| Hit16 | 751-69 | 2.19 | 0.67 | 1.38 | 1.41 | 0.76 | 6.17 | 2.38 | 3.98 | 0.35 | 3.22 | 2.46 |
| Hit17 | 752-21 | 1.1 | 0.21 | 0.51 | 0.61 | 0.45 | 0.64 | 0.56 | 0.42 | 0.09 | 0.43 | 0.24 |
| Hit18 | 754-31 | 1.04 | 0.61 | 1.94 | 1.20 | 0.68 | 0.15 | 0.1 | 0.19 | 0.13 | 0.14 | 0.04 |
| Hit19 | 757-90 | 1.67 | 0.48 | 1.17 | 1.11 | 0.60 | 0.48 | 0.44 | 0.51 | 0.20 | 0.41 | 0.14 |
| Hit20 | 758-29 | NA | 1.97 | 10.81 | 6.39 | 5.76 | 2.63 | 3.36 | 3.51 | 0.92 | 2.60 | 1.19 |

**Table S2**

IC_50_ values were obtained using ELISA and FRET assays for decompressed peptides that inhibit the interaction between TSLP and IL-7Rα/TSLPR.

| Decompression peptide hits | IC_50_（μM） | |
| --- | --- | --- |
|  | ELISA assay | TR-FRET assay |
| PL03202006 | 2.89 | 0.19 |
| PL03501006 | 8.02 | 0.6 |
| PL03502017 | 2.92 | 0.78 |
| PL03502026 | 1.19 | 0.86 |
| PL03202003 | 2.32 | 2.6 |
| PL03202034 | 0.98 | 2.41 |
| PL03202035 | 4.05 | 9.56 |
| PL03202036 | 6.64 | 17.63 |
| PL03202037 | 1.48 | 1.2 |
| PL03501038 | 6.4 | 5.03 |
| PL03502070 | 7.18 | 1.11 |

**Table S3**

Statistic of the plate uniformity for FLIPR screening.

| Data | agonist screening | | | antagonist screening | | |
| --- | --- | --- | --- | --- | --- | --- |
|  | Buffer control | ACTH control | peptide sample | Buffer control | ACTH control | peptide sample |
| RFU | 790.70 | 4683.70 | 232.42 | 95.36 | 3649.87 | 4276.44 |
| SD | 171.20 | 355.46 | 141.37 | 34.59 | 266.78 | 361.27 |
| Z'factor | 0.59 | |  | 0.60 | |  |
| Z factor |  | 0.79 | |  | | 0.72 |

**Table S4**

Identified 80-mer hits inhibiting the MC3R receptor. The inhibition rate was obtained at a single concentration of 2μM.

| Hits ID | Hits ID | primary inhibition rate | Secondary Test1 inhibition rate | Secondary Test2 inhibition rate | Secondary Test3 inhibition rate | Average inhibition rate | IC_50_（μM） | SD |
| --- | --- | --- | --- | --- | --- | --- | --- | --- |
| 80-mer peptide 1 | 500-93 | 64.90% | 73.50% | 53.10% | 50.00% | 60.38% | 0.28 | 0.07 |
| 80-mer peptide 2 | 619-70 | 74.10% | 75.20% | 45.20% | 44.00% | 59.63% | 0.60 | 0.18 |
| 80-mer peptide 3 | 623-10 | 57.20% | 76.90% | 35.50% | 44.90% | 53.63% | 0.54 | 0.06 |
| 80-mer peptide 4 | 431-65 | 52.00% | 77.10% | 40.10% | 42.80% | 53.00% | 0.41 | 0.04 |
| 80-mer peptide 5 | 601-92 | 46.60% | 72.60% | 45.30% | 40.90% | 51.35% | 0.59 | 0.16 |

**Table S5**

Identified decompressed peptides with inhibitory effects on the MC3R receptor. The inhibition rate was tested at a concentration of 2μM. 'NA' indicates that the IC_50_ was not tested.

| ID | Conformation | Length  (AA) | Inhibition test1 | Inhibition test2 | Average | SD | IC_50_  （μM） |
| --- | --- | --- | --- | --- | --- | --- | --- |
| PL00803002 | Linear | 40 | 72.50% | 63.00% | 67.75% | 6.72% | 0.31 |
| PL00803003 |  |  | 58.80% | 64.60% | 61.70% | 4.10% | 0.82 |
| PL00803004 |  |  | 93.67% | 89.01% | 91.34% | 3.30% | NA |
| PL00803005 |  |  | 55.30% | 47.00% | 51.15% | 5.87% | 0.94 |
| PL00803006 |  |  | 58.60% | 61.50% | 60.05% | 2.05% | 0.94 |
| PL00803007 |  |  | 90.48% | 92.13% | 91.31% | 1.17% | NA |
| PL00803032 |  |  | 71.28% | 69.08% | 70.18% | 1.56% | NA |
| PL00803033 |  |  | 74.93% | 81.28% | 78.11% | 4.49% | NA |
| PL00803036 |  |  | 51.10% | 52.60% | 51.85% | 1.06% | 0.55 |
| PL00803037 |  |  | 52.50% | 52.80% | 52.65% | 0.21% | 0.89 |
| PL00803038 |  |  | 51.00% | 52.83% | 51.92% | 1.29% | 1.03 |
| PL00803039 |  |  | 49.20% | 64.59% | 56.90% | 10.88% | 0.47 |
| PL00803040 |  |  | 59.70% | 63.10% | 61.40% | 2.40% | 0.79 |

**Table S6**

IC_50_ value and SD of the active peptides in the third and fourth decompression.

| Hits ID | Length（aa） | IC_50_（μM） | SD |
| --- | --- | --- | --- |
| PL00803107 | 19 | 0.35 | 0.02 |
| PL00803108 | 18 | 0.68 | 0.05 |
| PL00803109 | 17 | 0.72 | 0.04 |
| PL00803110 | 16 | 0.59 | 0.01 |
| PL00803111 | 15 | 0.64 | 0.02 |
| PL00803112 | 14 | 0.26 | 0.08 |
| PL00803113 | 13 | 0.54 | 0.11 |
| PL00803134 | 12 | 1.50 | 0.12 |
| PL00803135 | 11 | 0.70 | 0.02 |
| PL00803136 | 10 | 0.51 | 0.06 |
| PL00803137 | 9 | 1.14 | 0.11 |
| PL00803138 | 8 | 9.49 | 1.56 |
| PL00803139 | 7 | 6.66 | 0.84 |
| SHU9119 | 7 | 0.50 | 0.16 |

**Table S7** Comparison of the IC_50_ value of hits on the inhibitory activation against MC3R and MC4R receptors.

| Hits ID | ACTH-induced MC3R cell activation | | NDP-α-MSH-induced cells MC3R activation | | NDP-α-MSH -induced MC4R cells activation |
| --- | --- | --- | --- | --- | --- |
|  | IC_50_（μM） | SD | IC_50_（μM） | SD | IC_50_（μM） |
| PL00803107 | 0.32 | 0.01 | 1.01 | 0.17 | 54.29 |
| PL00803112 | 0.04 | 0.00 | 0.05 | 0.00 | 79.55 |
| PL00803136 | 0.79 | 0.14 | 2.92 | 0.29 | 40.62 |
| SHU9119 | 0.08 | 0.00 | 0.16 | 0.04 | 0.14 |
